# Isolation and characterization of mollicute symbionts from a fungus-growing ant reveals genome reduction and host specialization

**DOI:** 10.1101/2024.08.02.606451

**Authors:** Emily A. Green, Ian Klepacki, Jonathan L. Klassen

## Abstract

Two mollicute species belonging to the *Mesoplasma* and *Spiroplasma* genera have been detected in several species of fungus-growing ants using molecular methods. However, their ecological roles remain largely only inferred from metagenomic data. To better understand their diversity and specialization, we cultured both of these *Mesoplasma* and *Spiroplasma* symbionts from the fungus-growing ant *Trachymyrmex septentrionalis*, providing the first isolated mollicutes from any fungus-growing ant species. The genomes of our isolates and related metagenome-assembled genomes (MAGs) from *T. septentrionalis* fungus gardens comprise two unique phylogenetic lineages compared to previously described *Mesoplasma* and *Spiroplasma* species, and from related MAGs previously sequenced from the leaf-cutting ant *Acromyrmex echinatior*. This suggests that the *T. septentrionalis* symbionts comprise undescribed species with distinct host specificities. *Mesoplasma* genomes and MAGs also demonstrate regional specificity with their *T. septentrionalis* ant hosts. Both *Mesoplasma* and *Spiroplasma* strains from *T. septentrionalis* can catabolize glucose and fructose; both sugars are common in the ant’s diet. Similarly, both these *Mesoplasma* and *Spiroplasma* can catabolize arginine but only *Mesoplasma* can catabolize *N*-acetylglucosamine, which could both produce ammonia for the ants or fungus garden. Based on our genomic and phenotypic analyses, we describe these *T. septentrionalis* symbionts as *Mesoplasma whartonense sp. nov.* and *Spiroplasma attinicola sp. nov.*, providing insight into their genomic and phenotypic diversity, and cultures to facilitate future studies of these common but poorly understood members of the fungus-growing ant symbiosis.

## INTRODUCTION

Symbiosis is defined as two or more organisms living together in close association [1]. Bacterial symbionts have many different types of relationships with such hosts (e.g., pathogenic, commensal, or mutualistic) [2]. These relationships can result in tissue damage [3], manipulation of reproduction [4], nutrient provision [5], or protection [6] of the host. Symbionts can be transmitted by vertical transmission (from mother to offspring), horizontal transmission (from the environment or an individual of the same generation), or a mixture of both [7]. Bacterial symbionts often undergo genome reduction, evolving to have smaller genomes by eliminating genes that are not needed by these symbionts [8, 9]. Reduced genomes can prevent such symbionts from making amino acids or replicating outside of their host, thus ultimately tightening the bacteria-host symbiosis.

One well-known class of genome-reduced bacterial symbionts are the Mollicutes. Mollicutes are rapidly evolving via high mutation rates and frequent genome rearrangements, and have reduced genome sizes (580–2,200 kbps) and low %GC contents compared to most other bacterial classes [10]. They lack cell walls and stain Gram-negative despite evolving from a Gram-positive lineage. Mollicutes are strictly host-associated and include the well-studied genera *Mycoplasma*, *Mesoplasma*, *Entomoplasma*, *Spiroplasma*, and many others [11]. *Mesoplasma* species grow in serum-free media, do not require cholesterol, and commonly ferment glucose. They have been isolated from many different plants and insects but have no known pathogenicity or benefit towards these hosts [11]. *Spiroplasma* species are better-studied and are associated with diverse plants, insects, crustaceans, and mammals [11, 12]. They can ferment glucose, and require serum but not sterols for growth in culture. Some *Spiroplasma* species are pathogens that infect citrus and corn plants, causing citrus stubborn disease and corn shunt disease, respectively [13, 14]. Others manipulate sex ratios in *Drosophila* or cause tissue damage in mosquitos [15, 16]. *Spiroplasma* can also be mutualistic, as in *Drosophila neotestacea* where they defend against nematode infections [17].

Recent work has described the common presence of Mollicute bacteria belonging to the order Entomoplasmatales (which contains both *Mesoplasma* and *Spiroplasma*) in fungus-growing ants (Formicidae: tribe Attini). These ants evolved 55 to 65 MYA in South America [18–20] and today include 19 genera and approximately 250 known species that range from Northern U.S.A to Southern Argentina [20–24]. Many of these ants farm a coevolved fungus from the genus *Leucoagaricus* as their obligate food source [18, 22, 23]. The ants forage materials for this fungus, including leaves, dried plant material, and insect frass [25], and the fungus produces hyphal swellings called gonglydia that the ants eat [26, 27]. Except for the defensive actinobacterium *Pseudonocardia* and the fungal pathogen *Escovopsis* [28, 29], other members of the symbiosis remain poorly studied.

Recent environmental 16S rRNA gene sequencing studies of ant guts have revealed the frequent and abundant presence of *Mesoplasma* and *Spiroplasma* in several Attine ant genera [30–34]. An amplicon sequencing variant (ASV; a unique sequence variant obtained by clustering environmental 16S rRNA gene sequencing datasets), named EntAcro1 and phylogenetically related to the genus *Mesoplasma*, was detected in worker ants, larvae and pupae of *Atta cephalotes* and *Acromyrmex echinatior* [32], and in *At. colombica*, *At. sexdens. Ac. octospinosus*, and *Paratrachymyrmex* (previously *Trachymyrmex*) *cornetzi* worker ants [33]. A second ASV, named EntAcro10 and phylogenetically related to the genus *Spiroplasma,* was detected in worker ants and pupae from *At. cephalotes*, and in worker ants from *At. colombica*, *At. sexdens*. *Ac. octospinosus, P. cornetzi*, *Sericomyrmex amabilis*, *Mycetomoellerius* (previously *Trachymyrmex*) *zeteki*, *Mycocepurus smithii*, *Cyphomyrmex longiscapus*, and *Apterostigma dentigerum*. EntAcro1 was more abundant than EntAcro10 in *At. cephalotes*, *At. colombica*, *At. sexdens*. *Ac. octospinosus*, and *P. cornetzi*, and the opposite was true for *S. amabilis* and *M. zeteki*; these differences were hypothesized to be due to differences in foraging between ant species [33]. The abundances of EntAcro1 and EntAcro10 differed between ants, and not every ant in a colony was colonized by either species. 16S rRNA gene ASVs that were related to both of these *Mesoplasma* and *Spiroplasma* and lineages were also detected in most *T. septentrionalis* ant colonies [30, 35]. However, only one of *Mesoplasma* or *Spiroplasma* was present in an individual *T. septentrionalis* colony and their abundances varied between individual ants [35]. The abundances of these ASVs also varied between ant castes and following adaptation of ant colonies to the laboratory environment [35]. Although these studies demonstrate the symbiotic relationship between Mollicutes and different genera of fungus-growing ants, these 16S rRNA gene datasets lack the phylogenetic resolution necessary to provide information concerning the specificity and evolutionary history of *Mesoplasma* and *Spiroplasma* in the fungus-growing ant symbiosis and are unable to reveal the ecological consequences of their relationships with their ant hosts.

It is unknown how mollicute bacteria are acquired by their ant hosts. Potentially, ants might acquire them from a natural vector (e.g., plants) or from another ant colony [36]. Neither *Mesoplasma* or *Spiroplasma* are likely to be transferred vertically between ant generations, because fluorescence *in situ* hybridization imaging showed that EntAcro1 and EntAcro10 colonized the fat bodies, midgut, Malpighian tubules, ileum, and rectum of *At. cephalotes*, *At*. *colombica*, *Ac. echinatior*, *Ac. octospinosus*, *M. zeteki*, and *S. amabilis*, the fat bodies and ileum of *P. cornetzi,* and the midgut, ileum and rectum of *C. longiscapus* [32, 33]. Neither EntAcro1 or EntAcro10 was found in the reproductive organs of any of these ants, or either in or on ant eggs. Instead, mollicute bacteria are most likely socially transmitted from worker to worker, as demonstrated by the colonization of *Ac. echinatior* callows (freshly emerged ants) only when they were allowed to interact with older, colonized, worker ants [32].

Metagenomic sequencing can provide a semi-complete genome of microbes that are difficult to culture in isolation. Metagenomic assembled genomes (MAGs) can be computationally reconstructed from such data to generate hypotheses about a microbe’s potential function, but microbial isolates are needed to test such hypotheses. Although MAGs provide more functional insight than the 16S rRNA gene, a MAG may not comprise a complete bacterial genome [37, 38], and the expense of metagenomic sequencing can limit the number of unique MAGs available to study an individual bacterial species.

Metagenome-assembled genomes have been generated from a single *Ac. echinatior* metagenome for both *Mesoplasma* EntAcro1 and *Spiroplasma* EntAcro10 [39]. The MAGs for both *Mesoplasma* EntAcro1 and *Spiroplasma* EntAcro10 included genes predicted to decompose excess arginine into ammonium, which were hypothesized to be provided to the fungus garden via ant fecal droplets [39]. *Mesoplasma* EntAcro1 was also predicted to catabolize citrate, which might be acquired from fruit, leaves, or other plant material foraged by the ants, transforming it into acetate that might then be secreted by the bacterium and imported by the ants. Based on these data, Sapountzis et al. [39] hypothesized that EntAcro1 and EntAcro10 are nutritional mutualists that use these citrate and arginine catabolic pathways to provide nutrients for the ants and fungus garden. However, these hypotheses have yet to be experimentally tested, especially using cultured bacteria.

To overcome the limitations of the sequence-based techniques that have thus far been used to characterize mollicute symbionts of fungus-growing ants, in this study we isolated several *Mesoplasma* and *Spiroplasma* symbionts from *T. septentrionalis* ant guts. Our genomic comparisons using these isolates and related MAGs assembled from multiple *T. septentrionalis* fungus garden metagenomes [40] shows how *T. septentrionalis* symbiont genomes differ in size, gene content, and phylogenetic position from the EntAcro1 and EntAcro10 MAGs isolated from *Ac. echinatior*, suggesting that distinct phylogenetic lineages of Mollicutes are specific to different fungus-growing ant hosts. We also discovered that *Mesoplasma* symbionts from *T. septentrionalis* form phylogenetic clusters that are each unique to specific geographic regions. The results of our phenotypic tests partially validated the predictions that we made based on our gene annotations. Together, these results reveal a highly specific relationship between *T. septentrionalis* ants and their mollicute symbionts and set the stage for future experiments that move beyond exclusively culture-independent approaches to experimentally confirm the effects of these Mollicutes on the other members of the fungus-growing ant symbiosis.

## METHODS

### Sample Collection

*Trachymyrmex septentrionalis* colonies were collected from New Jersey, North Carolina, Georgia, and Florida between 2013 and 2020 (Supplemental Table 1). Permits for collection include New Jersey Department of Environmental Protection Division of Parks and Forestry State Park Service unnumbered letters of authorization, North Carolina Division of Parks and Recreation scientific collection and research permit 2015_0030, Georgia Department of Natural Resources State Parks and Historic Sites scientific research and collection permit 032015, and Florida Department of Agriculture and Consumer Services unnumbered letters of authorization. Directly after colony collections, fungus garden samples were stored in DESS (20% dimethyl sulfoxide, 250 mM disodium EDTA, and saturated sodium chloride) on dry ice [41]. The remainder of these ant colonies were transported to the University of Connecticut (U.S. Department of Agriculture permit P526P-14-00684) and maintained in a temperature-controlled room. Similar to [42], colonies were housed in 6 ¾ in x 4 13/16 in x 2 3/8 in plastic boxes lined with plaster of Paris, watered biweekly to maintain near-100% humidity, and provided sterile cornmeal *ad libitum* as food for the fungus garden. Frozen fungus garden samples were stored at −80 ℃ until DNA extraction.

### Isolation & Growth

Live ants were taken from the lab colonies, submerged in ethanol for 10 seconds, then submerged again in phosphate buffered saline (PBS) for 10 seconds; washes were repeated three times per ant. The gut (hindgut, midgut and rectum) was dissected from each ant abdomen using Swiss Style Superfine Tip 112 mm forceps (BioQuip, Rancho Dominguez, CA), placed into a 2 mL screw top tube filled with 1 mL PBS, and ground using a small sterile pestle. Cultures were grown in SP-4 modified from [43] as follows: Difco PPLO Broth Base (without crystal violet) replaced the Mycoplasma Broth Base, and both 20 mg of phenol red and 630 mL of water was added to the basal medium before autoclaving. The supplements used were 200 mL of fetal bovine serum, 25 mL of CMRL 1066, 5 mL of 1 N NaOH, and 5 mL of a 40% L-arginine solution (replacing α-ketoglutaric acid). No penicillin was added at this stage. These supplements were sterilized using a 0.45 µm filter before being added to the autoclaved basal medium. One hundred microliters of homogenized ant gut solution was added to 15 mL Falcon tubes containing either: 1) 4 mL of modified SP-4 broth, 2) 2 mL of modified SP-4, or 3) 4 mL modified SP-4 containing 40 mg/mL penicillin (Fisher Scientific, Hampton NH). Tubes 1 and 3 were incubated at 30 °C. Tube 2 was incubated at 37 °C for 1 hour, filtered using a 0.45 µM 30 mm polypropylene membrane syringe filter into 2 mL of fresh modified SP-4 resulting in 4 mL final volume, and incubated at 30 °C. Cultures were removed from the incubator after they changed color from red to orange/yellow or became turbid, typically within 5 to 14 days. Cultures that remained red with no turbidity were removed after four weeks. We purchased *Mesoplasma lactucae* 831-C4 (ATCC 49193) and *Spiroplasma platyhelix* PALS-1 (ATCC 51748) as reference strains. The freeze-dried pellets received from the ATCC were added to SP-4 broth and incubated at 30 °C until they changed color from red to orange/yellow. We described the sequencing of *S. platyhelix* ATCC 51748 previously [44].

### DNA Extraction & Sequencing

To choose cultures for purification, we performed community 16S rRNA sequencing using 1 mL of broth from each culture. DNA was extracted using the Epicentre MasterPure Complete DNA and RNA Purification Kit (Lucigen, Middleton WI) following the DNA purification protocol for cell samples and quantified using the Qubit dsDNA high-sensitivity assay protocol and a Qubit 3.0 fluorimeter (Invitrogen, Carlsbad CA). The presence of bacterial DNA was confirmed using PCR. Ten nanograms of template DNA was added to 5 µL Green GoTaq Reaction Mix Buffer (Promega, Madison, WI), 1.25 units of GoTaq DNA Polymerase (Promega, Madison, WI), 10 µmol of primers 515F and 806R (targeting the bacterial 16S rRNA gene V4 region) [45], and 300 ng bovine serum albumin (BSA; New England BioLabs Inc. Ipswitch, MA), to which nuclease-free H_2_O was added to a volume of 25 µL. Thermocycler conditions (BioRad, Hercules, CA) were: 3 min at 95 °C, 30 cycles of 30 sec at 95 °C, 30 sec at 50 °C, and 60 sec at 72 °C, followed by a 5 min cycle at 72 °C and then an indefinite hold at 4 °C. Gel electrophoresis confirmed the expected band size of 300–350 bp.

Samples were prepared for 16S rRNA community amplicon sequencing at the University of Connecticut Microbial Analysis, Resources and Services (MARS) facility. Approximately 30 ng of DNA from each sample was added to a 96-well plate containing 10 µmol each of the forward and reverse Illumina-barcoded versions of primers 515F and 806R, 5 µL AccuPrime buffer (Invitrogen, Carlsbad, California), 50 mM MgSO_4_ (Invitrogen, Carlsbad, California), 300 ng/µL BSA (New England BioLabs Inc. Ipswitch, Massachusetts), a 1 µmol spike-in of non-barcoded primers 515F and 806R, and 1 unit AccuPrime polymerase (Invitrogen, Carlsbad, California), to which nuclease-free H_2_O was added to a volume of 50 µL. Reaction mixes were separated into triplicate reactions (each with a volume of 16.7 µL) in a 384 well plate using an epMotion 5075 liquid handling robot (Eppendorf, Hamburg, Germany). This 384 well plate was transferred to a thermocycler (Eppendorf, Hamburg, Germany) that used the following conditions: 2 min at 95 °C, 30 cycles of 15 sec at 95 °C, 60 sec at 55 °C, and 60 sec at 68 °C, followed by a final extension for 5 min at 68 °C and then an indefinite hold at 4 °C. After PCR, triplicate reactions were re-pooled using the epMotion and DNA concentrations were quantified using a QIAxcel Advanced capillary electrophoresis system (QIAgen, Hilden, Germany). Samples with concentrations > 0.5 ng/µL were pooled using equal weights of DNA to create the final sequencing libraries. Libraries were then bead-cleaned using Mag-Bind RXNPure plus beads (OMEGA, Norcross, Georgia) in a 1:0.8 ratio of sequencing library to bead volume. Cleaned library pools were adjusted to a concentration of 1.1 ng/µL ± 0.1 ng/µL, which was confirmed using the Qubit dsDNA high-sensitivity assay on a Qubit 3.0 fluorimeter and sequenced as 2 x 250 bp libraries on an Illumina Miseq (Illumina, San Diego, California) at the UConn MARS facility. These data were deposited in the NCBI SRA database (SRR18059736-SRR18059804)

16S rRNA community amplicon sequencing reads were analyzed using R v3.5.3 [46] and the DADA2 v1.11.1 [47] pipeline for amplicon sequence variants (ASVs). Reads were classified using the SILVA database v128 [48, 49].

Cultures that contained *Spiroplasma* or *Mesoplasma* based on the community 16S rRNA gene sequencing, and the culture containing our reference *M. lactucae* strain, were T-streaked onto modified SP-4 agar plates, wrapped in parafilm to prevent the agar from drying, placed in a plastic bag or a candle jar (without a candle), and incubated at 30 °C. The identity and purity of each strain were confirmed by Sanger sequencing. Single colonies were taken from the agar plates and added to 5 µL Green GoTaq Reaction Mix Buffer (Promega, Madison, WI, USA), 1.25 units GoTaq DNA Polymerase (Promega, Madison, WI, USA), 10 μmol each of primers 27F and 1492R [50] (targeting the near-complete 16S rRNA gene), 300 ng/μL BSA (New England BioLabs Inc. Ipswitch, MA), and 50 mM MgCl_2,_ to which nuclease-free H_2_O was added for a final volume of 25 µL. Thermocycler conditions were: 2 min at 95 °C, 34 cycles of 1 min at 95 °C, 1 min at 54 °C and 1:30 min at 72 °C, a final extension for 5 min at 72 °C, and then a final hold at 4 °C. Gel electrophoresis confirmed the expected band size between 1000-1400 bp. These PCR products were bead cleaned using AMPure XP beads (Beckman Coulter, Brea, CA, USA), 8 µL of cleaned PCR product was added separately to 4 µL of primers 27F and 1492R, and the reactions were sent to Eurofins Genomics (Louisville, KY) for Sanger sequencing. Sanger sequencing chromatograms were analyzed using Geneious v2019.1.3 and their identities were confirmed using NCBI Nucleotide BLAST (https://blast.ncbi.nlm.nih.gov/, queried during

February, March, May, and July of 2021). These data were deposited in the NCBI GenBank database (OM760043-OM760044, OM812022-OM812029) For strains identified as *Mesoplasma* or *Spiroplasma*, a single colony from each source agar plate was spread over another SP-4 agar plate to increase cell concentrations. Genomic DNA was extracted from the resulting colonies using the Epicentre MasterPure Complete DNA and RNA purification kit, quantified, and confirmed as bacterial as described above. Genome sequencing libraries were then prepared using the Illumina TruSeq DNA PCR Free kit by the UConn MARS facility and sequenced on an Illumina MiSeq at the UConn Center for Genome Innovation as 2 x 250 bp libraries (Illumina, San Diego, California).

### Metagenome-assembled and reference genomes

Fungus garden bacterial metagenomes from *T. septentrionalis* colonies (which differed from those used for the Mollicute isolations) were sequenced and assembled by the Joint Genome Institute (JGI), who then generated metagenome-assembled genomes (MAGs) as described in [40]. The MAGs from these datasets with a Bin Lineage annotation of “Mollicutes” were downloaded from the JGI Integrated Microbial Genomes and Microbiomes website (https://img.jgi.doe.gov/). Additional MAGs (EntAcro1 and EntAcro10) that had been previously generated from *Ac. echinatior* ants [39] and other reference *Mesoplasma* and *Spiroplasma* genomes were downloaded from the NCBI (Supplementary Tables S2-S4).

### Genome Assembly, Quality checking, & Analysis

Sequencing adapters were removed using Trimmomatic v0.39 with default parameters [51], and the data were quality checked using FastQC v0.11.8 (https://www.bioinformatics.babraham.ac.uk/projects/fastqc/). Trimmed reads were assembled using unicycler v0.4.8-beta [52] and all genome assemblies (including those obtained from NCBI) were annotated using Prokka v1.13.3 [53]. Contigs < 2000 bp were checked for contamination using NCBI’s Nucleotide BLAST (https://blast.ncbi.nlm.nih.gov/) and investigating if taxonomic identity of their top BLAST hits matched to the publicly available *Mesoplasma lactucae* genome (NCBI accession GCA_002441935.1), MAG EntAcro1 [39], our published *Spiroplasma platyhelix* genome [44], or MAG EntAcro10 [39]. Genome completeness was checked using BUSCO v5.0.0 with the tenericutes_odb10 Tenericutes database [54]. The assembled genomes were deposited in the NCBI WGS (Supplementary Table S5) and the sequencing reads were deposited in the NCBI SRA database (SRR17695994-SRR17696005)

Prokka gene annotations for each genome were imported into anvi’o v7 [55, 56] using the gff.parser script (https://merenlab.org/2017/05/18/working-with-prokka/) and an anvi’o database was created for each genome following the anvi’o metagenome tutorial (https://merenlab.org/2016/06/22/anvio-tutorial-v2/). We followed the anvi’o pangenome and phylogenomics tutorials (https://merenlab.org/2016/11/08/pangenomics-v2/ and https://merenlab.org/2017/06/07/phylogenomics/) to calculate the genome length, %GC, and number of single-copy core gene (SCG) clusters (homologous genes present in all analyzed genomes), as well as dendrograms to show the presence/absence of gene clusters (i.e., homologous genes) in each genome. The anvi’o ‘--mcl-inflation’ parameter used to identify homologous genes can be set between 2 (less strict, used for distantly related genomes) and 10 (more strict, used for closely related genomes, typically of the same ‘species’ or ‘strain’). All analyses used an mcl inflation score of 7; results were the same using a score of 3 (Suppl. Fig. 1).

Assembled genomes were also functionally annotated using the RAST (Rapid annotation using subsystems technology) webserver [57] (https://rast.nmpdr.org/), selecting the Domain ‘Bacteria’, the appropriate genus (*Mesoplasma* or *Spiroplasma*), and ‘Genetic code 4’ as options. We used the ‘Classic RAST’ RAST annotation scheme, RAST as the gene caller, and FIGfam version ‘Release70’. RAST annotations are collated in Suppl. Table S6.

The anvi’o commands ‘anvi-get-sequences-for-gene-clusters’ and ‘--concatenate-gene-clusters’ were used to export a file that contained a concatenated alignment of all the amino acid sequences from the single-copy gene clusters, and the flag ‘--report-DNA-sequences’ was used to export their corresponding nucleotide sequences. Amino acid phylogenies were generated using FastTree (v.2.1.10) [58] with the Whelan-And-Goldman 2001 and CAT substitution model (-wag flag) and local bootstrapping. Nucleotide single-copy core gene cluster phylogenies were generated similarly but instead using the Jukes-Cantor model (-nt flag). Trees were visualized using iTOL (https://itol.embl.de/) [59]. Average nucleotide identity (ANI) values were computed in anvi’o using the ‘anvi-compute-genome-similarity’ command.

Five hundred and fifty “core” gene clusters, which include both single-copy core genes and other highly conserved core genes that were nearly-always present (but not absolutely so, e.g., due to incomplete genome and MAG assembly) were selected from our *Mesoplasma* genomes and MAGs, the EntAcro1 MAG, and the *M. lactucae* reference genomes using the anvi’o “Search gene clusters using filters” tab in the anvi’o interactive interface. The ‘minimum number of genomes gene cluster occurs’ was set to 18 and the ‘maximum functional homogeneity index’ was set to 0.99.The six hundred and eighty-four “core” gene clusters were similarly selected from our *Spiroplasma* genomes, the EntAcro10 MAG, and the *S. platyhelix* reference genome, setting the ‘minimum number of genomes gene cluster occurs’ to 3 and the ‘maximum functional homogeneity index’ to 0.99. The command ‘anvi-get-sequences-for-gene-clusters’ was used to export all the amino acid sequences from these core gene clusters, and the flag ‘--report-DNA-sequences’ was used to export their corresponding nucleotide sequences. ORFcor [60] and a custom perl script provided in the Supplemental Code was used to align and concatenate all these sequences into single .faa and .fna files for each genus. Phylogenies were created as described above.

### Phenotypic Tests

Metabolism of glucose, fructose, *N*-acetylglucosamine, urea and arginine were tested following the methods in [61], except using 0.04% (w/v) for chitin (which was otherwise insoluble). Sterol tests contained modified SP-4 lacking serum, modified SP-4 with tween and palmitic acid, modified SP-4 with tween, palmitic acid, and ethanol, and modified SP-4 containing 0, 1, 5, 10, and 20 µg/mL of cholesterol (MP Biomedicals), following [62] but using modified SP-4 medium instead of mycoplasma broth. Growth at 20, 25, 30 or 37 ℃ was tested in modified SP-4 medium without other supplements. Before inoculation, all strains were grown in 5 mL of modified SP-4 medium until an orange color became visible, indicating that the cells entered logarithmic growth phase, after which 100 µL was added to each phenotypic test. All cultures were incubated at 30 °C for four weeks or until they changed color from red to orange/yellow (indicating acidification) or from red to magenta (indicating alkalinization), typically within 5 to 14 days. *M. lactucae* ATCC 49193 and *S. platyhelix* ATCC 51748 were used as controls.

### Transmission electron microscopy imaging

Cultures of JKS002660 and JKS002670 were grown in 5 mL of modified SP-4 medium until an orange color became visible, indicating cells entering into logarithmic growth phase. These samples were then centrifuged at 5,000 rcf and the supernatant was removed. The cell pellets were resuspended in 1 mL of 0.1 M phosphate buffer, centrifuged at 5,000 rcf, and the phosphate buffer supernatant removed. This was repeated 3 times. Cells were then fixed in 2% glutaraldehyde (Electron Microscopy Sciences, Hatfield, Pennsylvania) for 1 hour and rinsed 3 times with 0.1 M phosphate buffer.

Imaging was conducted in the University of Connecticut’s Bioscience Electron Microscopy Laboratory using a FEI Tecnai 12 G2 Spirit BioTWIN transmission electron microscope. Three microliters of each cell suspension in 0.1 M phosphate buffer was added to a carbon copper mesh grid, to which 0.5% w/v uranium acetate (Electron Microscopy Sciences, Hatfield, Pennsylvania) was added to the grid and allowed to air dry.

## RESULTS

### Isolation

We dissected the guts of 69 *T. septentrionalis* ants from 17 colonies. Individual guts were crushed in PBS and inoculated into each of three enrichment conditions: (1) modified SP-4 medium; (2) modified SP-4 medium supplemented with penicillin; and (3) modified SP-4 medium that was heated and filtered (Suppl. Fig. S2). One hundred of the resulting 207 enrichment cultures became turbid or changed color (phenol red turns from red to orange/yellow due to acid production during fermentation; Suppl. Fig. S2). We characterized a subset of these cultures using 16S rRNA community amplicon sequencing (Suppl. Fig. S3) and inoculated samples containing *Spiroplasma* or *Mesoplasma* onto modified SP-4 agar plates. A single colony from each plate with a ‘fried egg’ morphology typical of Mollicutes was purified and taxonomically identified using 16S rRNA gene Sanger sequencing. Strains JKS002657 and JKS002669, JKS002658 and JKS002659, and JKS002663 and JKS002664 were isolated from different enrichment cultures inoculated using the same ant colonies (Suppl. Table S1); all other strains are from different ant colonies. In total, we isolated and identified 8 *Mesoplasma* and 3 *Spiroplasma* strains from 9 unique ant colonies (strains JKS002657-JKS002664 and JKS002669-JKS002671, respectively; Suppl. Tables S1 & S5).

### Gene Content

We sequenced the genomes of all 8 *Mesoplasma* and 3 *Spiroplasma* strains that we isolated from dissected *T. septentrionalis* guts. The 16S rRNA genes in these genomes were 100% identical to the amplicon sequence variants (ASVs) that we identified in the gut enrichment cultures (Suppl. Table S7) and to ASVs that we previously identified in the *T. septentrionalis* microbiome [35]. We also analyzed 11 *Mesoplasma* metagenome-assembled genomes (MAGs) that our lab sequenced in collaboration with the Joint Genome Institute (Suppl. Table S2) [40]. There were no *Spiroplasma* MAGs assembled from these metagenomes. For comparison, we also downloaded the MAGs assembled by Sapountzis et al. [39] (Suppl. Table S3) [39]. As controls, we also re-sequenced the genome *Mesoplasma lactucae* ATCC 49193 (which matched that sequenced previously; NCBI BioProject: PRJNA412357; Suppl. Table S4) and included our previously sequenced *Spiroplasma platyhelix* ATCC 51748 genome [44].

All 8 *Mesoplasma* strains that we isolated from *T. septentrionalis* guts had genome sizes of 762–785 kbps and GC contents of 32.68–32.79% (Suppl. Table S5). The *Mesoplasma* MAGs generated from *T. septentrionalis* fungus gardens [40] had lengths of 385–760 kbps and 32.41– 33.00% GC content. These MAGs were estimated to be 52.70–100% complete using BUSCO (Suppl. Table S2), with the smallest genomes being the least complete. The *Mesoplasma* EntAcro1 MAG from the fungus-growing ant *Ac. echinatior* had a genome size of 866 kbps and a GC content of 33.74% [39], which was larger in length and slightly higher in GC content compared to the *Mesoplasma* genomes and MAGs from *T. septentrionalis* (Suppl. Table S3). All of these genomes and MAGs were smaller than the *M. lactucae* ATCC 49193 genome (824 kbps), which also had a lower GC content (29.63%; Suppl. Table S4).

The genomes from all 3 *Spiroplasma* strains that we isolated from *T. septentrionalis* guts had sizes of 835–884 kbps and 25.49–25.77% GC content (Suppl. Table S5). The *Spiroplasma* EntAcro10 MAG from *Ac. echinatior* [39] had a similar genome size (838 kbp) and GC content (25.53%) to those of the *Spiroplasma* strains isolated from *T. septentrionalis* (Suppl. Table S3). The *S. platyhelix* ATCC 51748 genome was shorter than all of these genomes and MAGs (739 kbps) and its GC content was higher (28.60%; Suppl. Table S4).

The *Mesoplasma* genomes from the strains that we isolated contained 660–689 coding sequences. Similarly, the *T. septentrionalis* MAGs with 100% complete BUSCO scores contained 654–663 coding sequences. MAG EntAcro1 contained substantially more coding sequences (757) than the genomes and MAGs from *T. septentrionalis*. The *M. lactucae* reference genome contained 691 coding sequences, similar to the isolate genomes. Our pangenome analysis separated gene clusters from these *Mesoplasma* genomes and MAGs into three bins: (1) “fully conserved”, containing anvi’o “single-copy core genes”, i.e., single-copy genes present in all analyzed genomes; (2) “highly conserved”, genes found in most analyzed genomes; and (3) “variable”, genes found in only a few genomes. There were 57, 583, and 555 gene clusters in the fully conserved, highly conserved, and variable bins (Fig. 1A). Consistent with the “variable” genes being part of a poorly characterized accessory genome, 87.57% of these genes had no Prokka annotation, compared to 39.96%, and 35.08% of the “fully conserved” and “highly conserved” genes.

**Figure 1).**
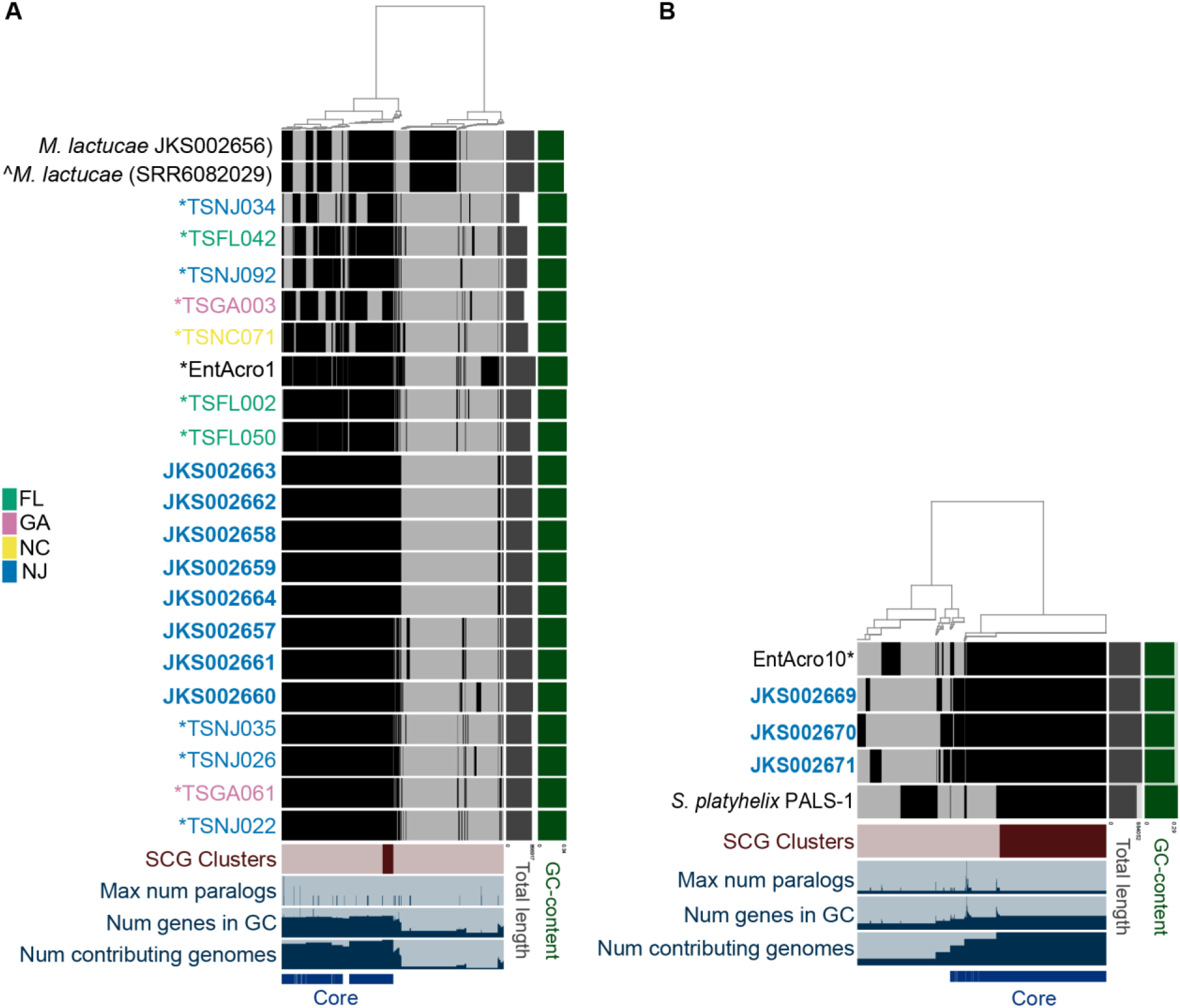
Anvi’o pangenome analysis of: A) *M. whartonense* genomes (bold) and MAGs (*), the EntoAcro1 MAG (*), and the *M. lactucae* genomes as a reference and B) *S. attinicola* genomes (bold), the EntAcro10 MAG (*), and the *S. platyhelix* genome as a reference. Text colors indicate strains or MAGs from the same state. Two *M. lactucae* genomes are shown, with the ^ indicating the genome downloaded from NCBI (SRR6082029) vs. the other that we re-sequenced as a control (JKS002656). In the central heatmap, black and grey bars indicate homologous genes that are present or absent in each genome. The dendrogram clusters groups of homologous genes (anvi’o gene clusters) using Euclidean distances.

The *Spiroplasma* genomes from the strains that we isolated contained 768–820 coding sequences, similar to MAG EntAcro10, which had 774 (Suppl. Table S6). The *S. platyhelix* reference genome contained significantly fewer coding sequences (670) than the *Spiroplasma* strains that we isolated. The *Spiroplasma* pangenome (Fig. 1B) had 1,111 gene clusters. There were 472, 156, and 483 gene clusters in the fully conserved, highly conserved, and variable bins with 26.81%, 80.77%, and 89.23% of these gene clusters having no gene annotation. The fewer *Spiroplasma* strains and lack of MAGs from *T. septentrionalis* compared to our *Mesoplasma* analysis likely caused the greater number of fully conserved genes and fewer highly conserved genes in our *Spiroplasma* analysis compared to that of *Mesoplasma*.

The *Mesoplasma* MAGs from *T. septentrionalis* fungus gardens closely resemble those of *T. septentrionalis* gut isolates, although those with low BUSCO scores lacked some ‘core’ genes, consistent with BUSCO predicting that these genomes are incomplete. Some small gene content differences occurred between related strains in both the *Mesoplasma* and *Spiroplasma* pangenomes, with the strains having the most similar gene content also clustering together in our phylogenetic trees (Fig. 2A & B, Suppl. Fig. S4-S9). Compared to the *T. septentrionalis* genomes and MAGs, the EntAcro MAGs lacked very few ‘core’ genes (i.e., those in the “fully conserved” and “highly conserved” gene bins). The reference genomes from *M. lactucae* and *S. platyhelix* had the most genes found only in their genome and lacked many of the ‘core’ genes found in the ant symbiont genomes and MAGs. Both *M. lactucae* genomes had 265 gene clusters that were found only in their genomes and lacked 34 *Mesoplasma* ‘core’ gene clusters. *S. platyhelix* had 158 gene clusters found only in its genome and lacked 22 *Spiroplasma* ‘core’ genes. The MAGs EntAcro1 and EntAcro10 had 86 and 84 unique gene clusters, respectively, that were not found in the reference genomes or the *T. septentrionalis* genomes and MAGs. All genes unique to these reference genomes and MAGs were included in the ‘variable’ bins of our pangenome analyses.

**Figure 2).**
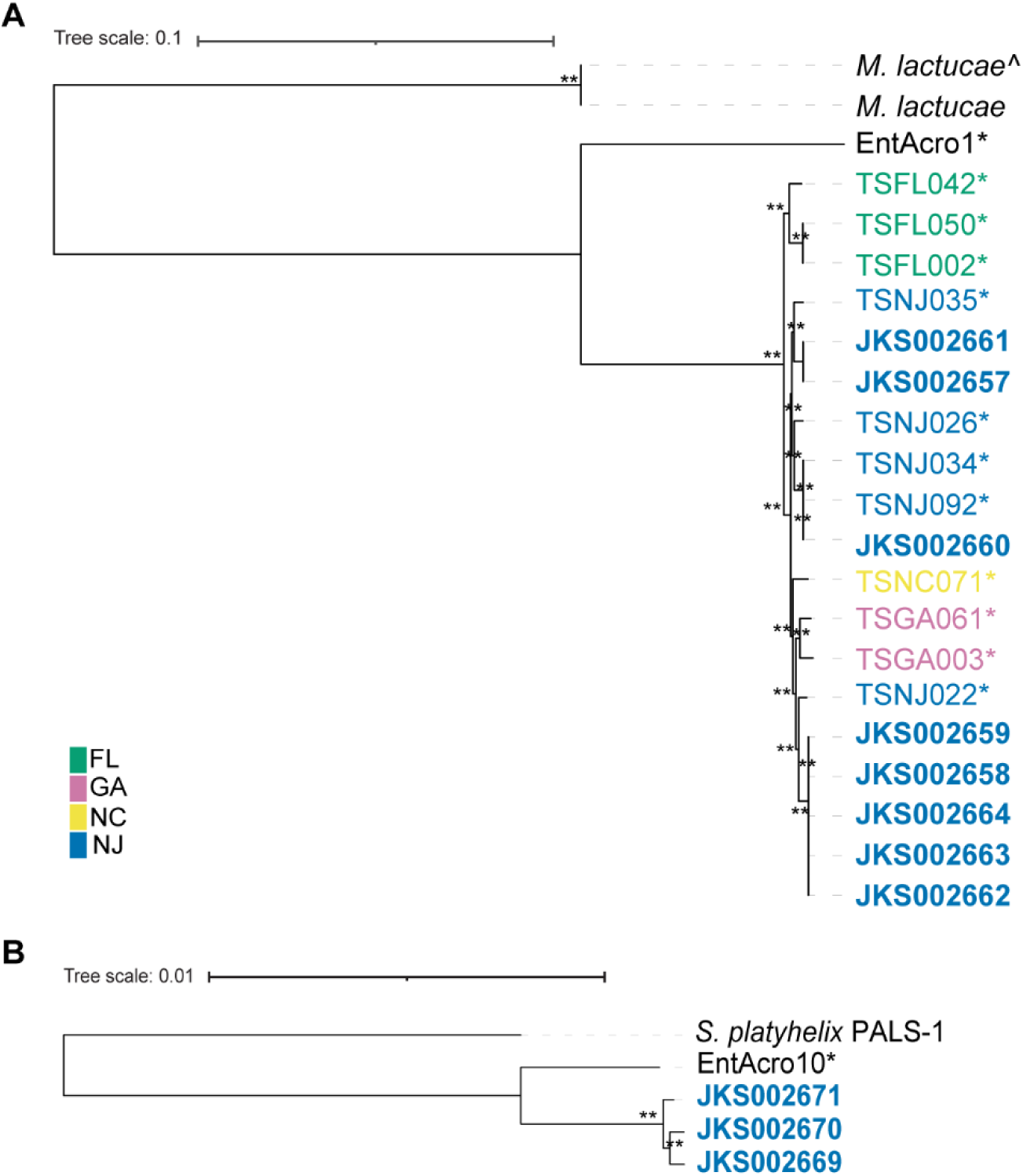
A) *Mesoplasma* nucleotide phylogeny created using 550 core gene clusters as input data and the fasttree JC + CAT substitution model. The tree was rooted using the midpoint of the branch leading to the *M. lactucae* genomes. The ^ indicates the NCBI (SRR6082029) genome. B) *Spiroplasma* nucleotide phylogeny created using 684 core gene clusters as input data and the fastree JC + CAT substitution model. The tree was rooted using the midpoint of the branch leading to the *S. platyhelix* genome. Asterisks indicate MAGs and bolded names indicate *T. septentrionalis* isolate genomes. Colors group strains by state. Local bootstrap values of 60–79 and 80–100 are indicated by * and **, respectively.

### Phylogeny & Average Nucleotide Identity

We created phylogenies using both amino acids and nucleotides from 21 single-copy gene clusters that were present in all ant-related *Mesoplasma* genomes and MAGs, as well as in several NCBI *Mesoplasma* reference genomes (Suppl. Fig. S4 & S5). In both the amino acid and nucleotide phylogenies, the NCBI reference *Mesoplasma* genomes grouped with a similar topology to the tree in Gasparich 2019 [63], in which *M. lactucae* forms an outgroup to all other *Mesoplasma* strains. All *Mesoplasma* ant-related genomes and MAGs formed their own clade next to *M. lactucae*. Similarly, we created phylogenies using all 80 single-copy core gene clusters that were present in all *Spiroplasma* genomes and MAGs from both ants and NCBI, using both amino acid and nucleotide sequences (Suppl. Fig. S6 & S7). Here, *S. platyhelix* formed an outgroup from the other NCBI *Spiroplasma* genomes, as in the *Spiroplasma* phylogenetic tree in the Bergey’s Manual of Systematics of Archaea and Bacteria [11], and all *Spiroplasma* genomes from ants formed their own clade next to *S. platyhelix*. Thus, both our *Mesoplasma* and *Spiroplasma* isolate genomes formed monophyletic clades close to but distinct from the reference species *M. lactucae* and *S. platyhelix*, respectively. We note that a previous proposal exists to reclassify these genera [64] but has not yet been formally adopted. We have therefore maintained the currently accepted names *Mesoplasma* and *Spiroplasma* in this study while acknowledging the potential need for future reclassification.

To increase the phylogenetic resolution of our analyses, we created additional phylogenies using the core gene clusters (i.e., those in the “fully conserved” and “highly conserved” bins) in all our ant genomes, all ant MAGs, and their closest reference genomes. The pangenome including only the *M. lactucae* reference genome, MAG EntAcro1, and the *Mesoplasma* isolate genomes and MAGs from *T. septentrionalis* had 550 core gene clusters that were used for these phylogenies. The *Spiroplasma* pangenomes including only the *S. platyhelix* reference genome, MAG EntAcro10, and the *Spiroplasma* isolate genomes from *T. septentrionalis* had 684 core gene clusters that were used for these phylogenies.

In both the amino acid and DNA sequence-based trees, the EntAcro MAGs from *A. echinatior* grouped separately from the rest of the *T. septentrionalis* genomes and MAGs (Figs. 2A & B, Suppl. Figs. S8 & S9), with the *M. lactucae* or *S. platyhelix* references genomes forming outgroups. In all *Mesoplasma* phylogenies, the *T. septentrionalis* genomes and MAGs clustered by the geographic locations from which their corresponding ant colonies had been collected (Fig. 2A. Suppl. Fig. S8). For example, all the *Mesoplasma* strains that we isolated were from ant colonies collected in New Jersey, and these genomes clustered together with New Jersey MAGs. North Carolina MAG TSNC071 formed its own branch in the DNA tree (Fig. 2A) but clustered with the New Jersey genomes and MAGs in the amino acid tree (Suppl. Fig. S8). The Florida and Georgia MAGs formed their own clades in both the amino acid and DNA sequence trees. This suggests that *T. septentrionalis Mesoplasma* strains are specific to distinct geographic locations. Because we lack *Spiroplasma* genomes and MAGs sampled from across a similarly broad geographic range, whether their genomes show a similar biogeographic trend remains unknown.

Average Nucleotide Identities (ANIs) confirmed that all *T. septentrionalis* genomes and MAGs are distinct from the NCBI genomes and EntAcro MAGs (Fig. 3A & B). All *Mesoplasma* genomes and MAGs from *T. septentrionalis* had 72–73% ANI to the *M. lactucae* reference genomes and 70–73% ANI to the other NCBI reference genomes. Similarly, the *Spiroplasma* genomes from *T. septentrionalis* had 78% ANI to the *S. platyhelix* reference genome and 70– 78% ANI to the other NCBI reference genomes. These results confirm that the *T. septentrionalis* genomes and MAGs should not be classified as these species [65, 66]. The *Mesoplasma* genomes and MAGs from *T. septentrionalis* had 87% ANI to the EntAcro1 MAG (Fig. 3A), and the *Spiroplasma* genomes from *T. septentrionalis* had 92% ANI to EntAcro10 (Fig. 3B). All *Mesoplasma* genomes and MAGs from *T. septentrionalis* had 99–100% similarity to each other, and these ANIs were not differentiated by state as in the phylogenies (Fig. 2A & Suppl. Fig. S8). Similarly, the *Spiroplasma* genomes from *T. septentrionalis* had ANIs of 98–100% when compared to each other (Fig. 3B). These values are consistent with all *T. septentrionalis Mesoplasma* and *Spiroplasma* belonging to their own species that are distinct from the EntAcro MAGs [65, 66]. We therefore named them *Mesoplasma whartonense sp. nov.* and *Spiroplasma attinicola sp. nov* and characterized them further below.

**Figure 3).**
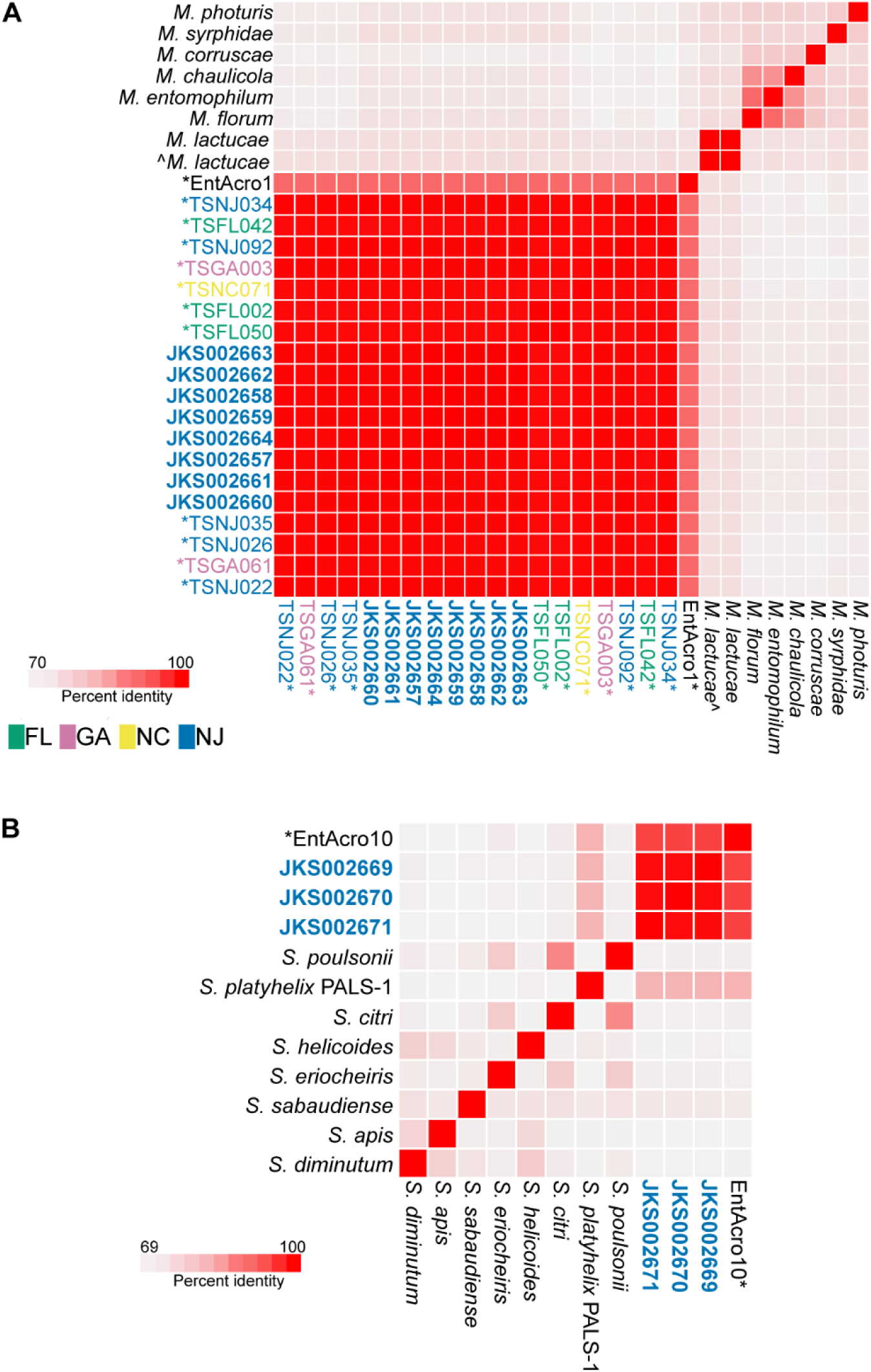
Average nucleotide identity heatmap of A) *M. whartonense* genomes and MAGs, MAG EntAcro1, and *Mesoplasma* reference genomes, and B) *S. attinicola* genomes, MAG EntAcro10, and *Spiroplasma* reference genomes, produced by anvi’o. Asterisks indicate MAGs bold names indicate our isolate genomes. The ^ indicates the NCBI (SRR6082029) genome and colors indicate the state from which strains or MAGs were collected.

### Gene functions

We used RAST to predict the functions of all genes in *M. whartonense* and *S. attinicola* (Fig. 4, Suppl. File 1). As expected for Mollicutes, *M. whartonense* and *S. attinicola* lacked the genes necessary to produce a cell wall. Only *M. whartonense* had genes for the biosynthesis of glycerophospholipids, but both *M. whartonense* and *S. attinicola* had genes for lipases that likely facilitate the acquisition of host-derived fatty acids for cell membrane biogenesis. Both species lacked the genes needed to produce many amino acids but retained genes for nucleotide salvage, suggesting they also likely depend on their ant hosts to provide many amino acids. Both *M. whartonense* and *S. attinicola* had the genes necessary to import vitamin B, folate, and riboflavin, but only *M. whartonense* had thiamine utilization genes.

**Figure 4).**
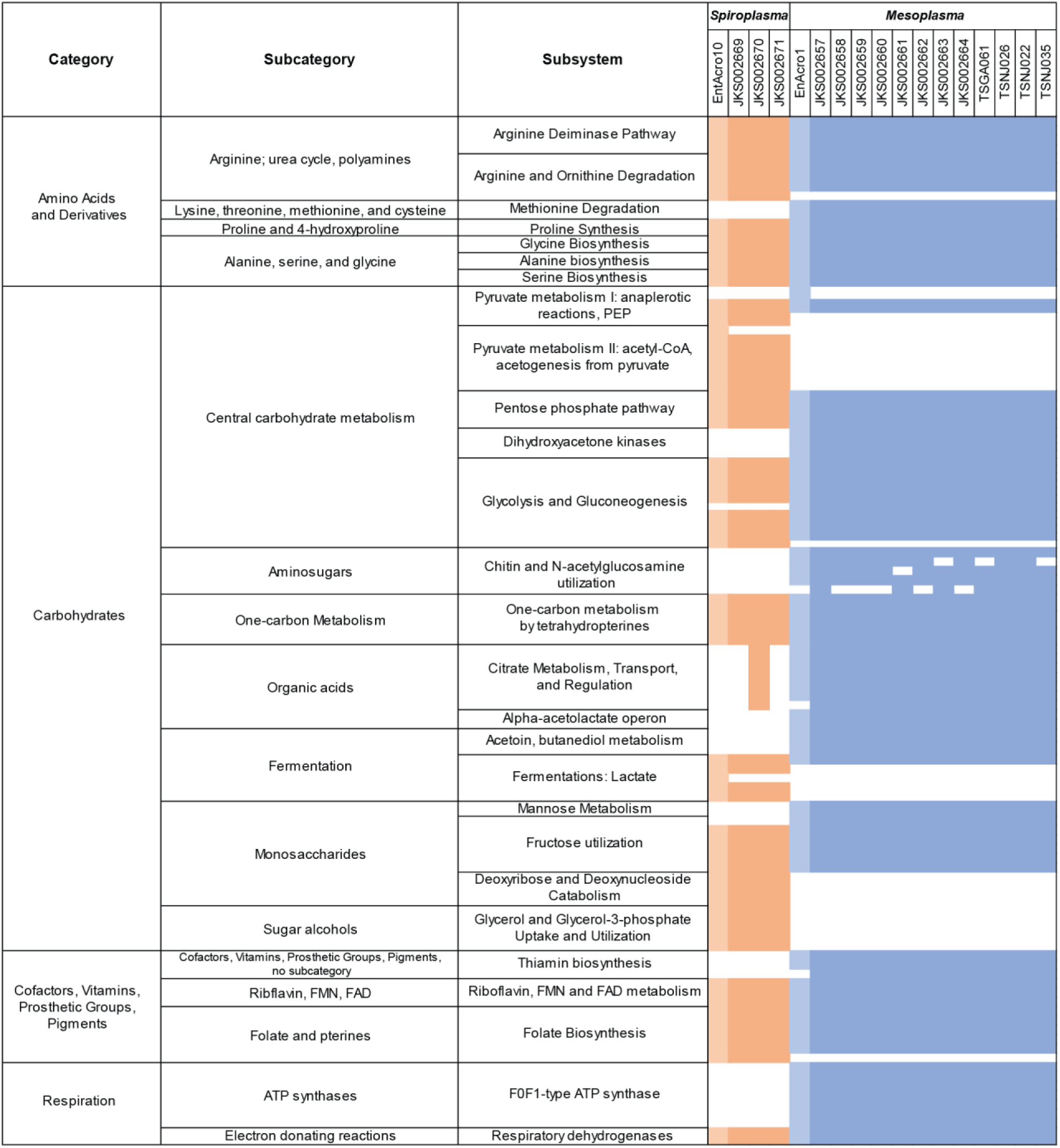
Representative gene function predictions for ant-associated *Spiroplasma* and *Mesoplasma*, as predicted using RAST. Orange colors indicate genes present for *Spiroplasma* and blue indicates genes present for *Mesoplasma*. The lighter of each color denotes genes present in MAGs EntAcro1 or EntAcro10. White indicates no presence.

Both *M. whartonense* and *S. attinicola* had genes encoding for bacteriophage defense systems, including a Type I restriction-modification system and CRISPR-Cas1, but neither had any obvious prophages. Only *S. attinicola* had genes for a Type III restriction-modification system, and only *M. whartonense* had genes for CRISPR-Cas2 (Fig. 4, Suppl. File 1). The conserved presence of such genes is consistent with both *M. whartonense* and *S. attinicola* being frequently exposed to bacteriophages, as also proposed for EntAcro1 and EntAcro10 [39].

Reconstructions of catabolism in *M. whartonense* and *S. attinicola* are shown in Fig. 5. We predict that both *M. whartonense* and *S. attinicola* can catabolize glucose and fructose using glycolysis and the pentose phosphate pathway to produce pyruvate, ATP, and NAD(P)H, and catabolize arginine to produce ornithine, ATP, and ammonium (NH_4_^+^); these catabolic pathways are present in many Mollicutes (Fig. 4, Suppl. File 1) [67]. Certain low-quality assemblies (especially the MAGs) lacked some genes present in their close relatives, but such differences are more likely technical than biological. Glucose and fructose likely originate from plant material that the ants forage or are produced during fungal digestion. The use of such digestion products is also suggested by *M. whartonense* having genes for chitin degradation and *N-* acetylglucosamine. Catabolism of sugars could produce pyruvate from the glycolysis or pentose phosphate pathways or provide precursors for the biosynthesis of purine and pyrimidines. We predict that both *M. whartonense* and *S. attinicola* possess fermentation pathways leading from pyruvate to L-lactate (both species), D-lactate (*S. attinicola*), acetoin (*M. whartonense*), and acetate (*S. attinicola*).

**Figure 5.**
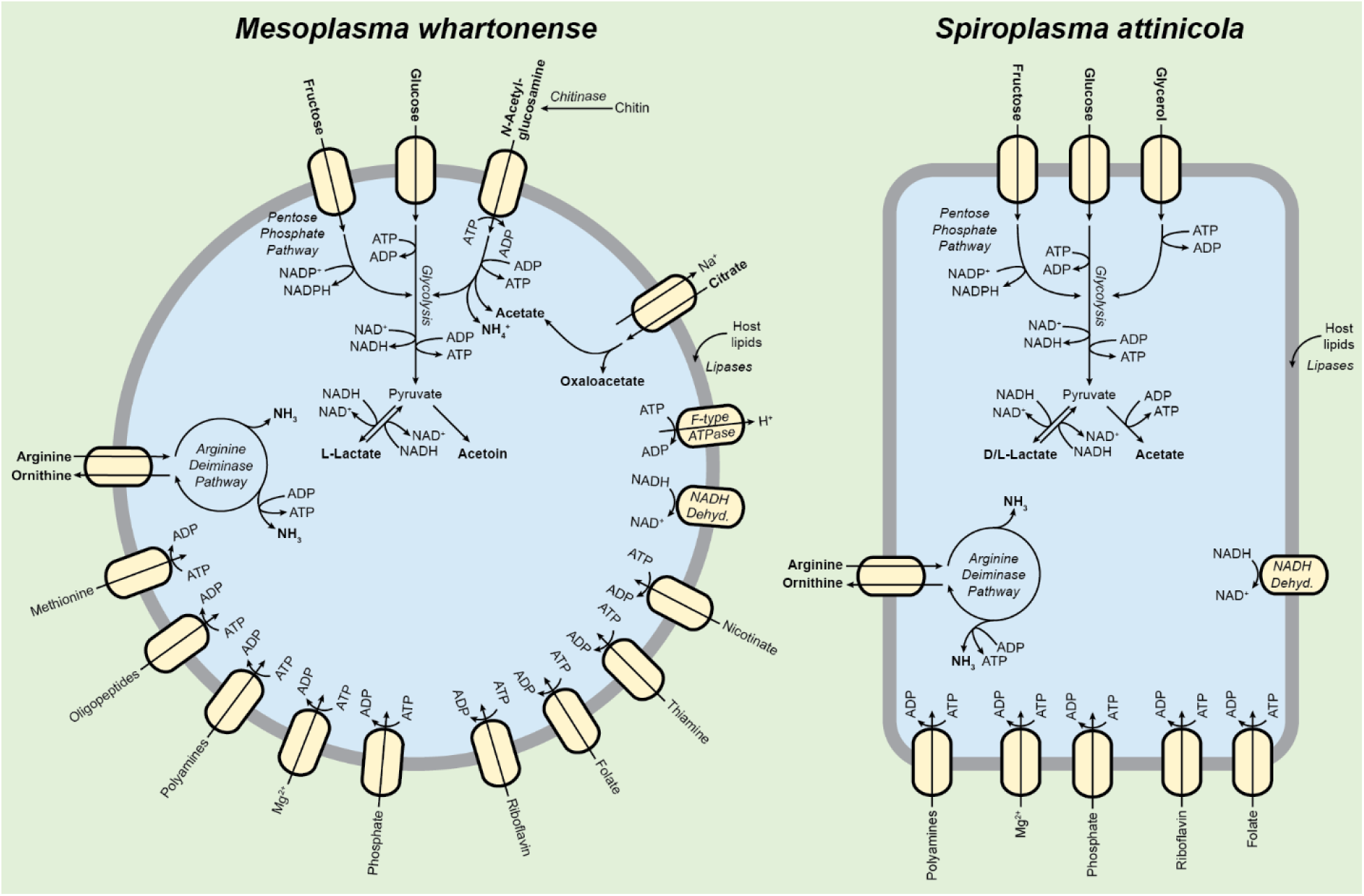
Representative catabolic pathways in *M. whartonense* (left) and *S. attinicola* (right). Anabolic reactions are not shown and reactions are not balanced. Major predicted catabolic substrates and products are bolded.

We predict that both *M. whartonense* and *S. attinicola* can catabolize dihydroxyacetone phosphate, but only *M. whartonense* can import glycerol. These are likely used to produce glyceraldehyde-3-phosphate that can enter the glycolysis pathway. Glycerol might also be produced by the lipases encoded by both *M. whartonense* and *S. attinicola* and may be used for glycerophospholipid biosynthesis in *M. whartonense* (but not *S. attinicola*, which lacks the corresponding genes). *M. whartonense* can also likely produce acetate, but from citrate (which could come from foraging materials) instead of from pyruvate as in *S. attinicola* (except for *S. attinicola* JKS002670, which we predict can produce acetate from both pyruvate and citrate). This acetate may be assimilated by the ant host [39]. Both *M. whartonense* and *S. attinicola* can likely interconvert glycine and serine, convert aspartate into fumarate, import and metabolize polyamines, and encode an NADH oxidase. Only *M. whartonense* has genes for the import and metabolism of methionine, and F0F1 ATPase genes that are predicted to mainly hydrolyze ATP to maintain an electrochemical gradient in other Mollicutes [68, 69].

The EntAcro1 MAG and the *M. whartonense* genomes had very similar gene contents, except for a few differences. The EntAcro1 MAG lacked genes for methionine transport, a Apo-citrate lyase phosphoribosyl-dephospho-CoA transferase, CRISPR-Cas2, and thiamine transport protein ThiT, which were all present in *M. whartonense* (Fig. 4; Supplementary File 1). Whether these absences are due to incomplete MAG assembly is unknown. In contrast, the EntAcro1 MAG did contain an NADP-dependent malic enzyme gene and a TrkH potassium uptake gene that were lacking in *M. whartonense*. The EntAcro10 MAG and *S. attinicola* genomes were more similar in length and gene count compared to the *Mesoplasma* genomes (Fig. 4). The EntAcro10 MAG lacked genes for a DNA-cytosine methyltransferase and the TolA protein, which were both present in *S. attinicola*. However, the EntAcro10 MAG contained several genes lacking in *S. attinicola*, such as an acetaldehyde dehydrogenase gene, two coenzyme A biosynthesis genes, a methylglyoxal metabolism gene and an internalin-like protein gene (Supplementary File 1). Overall, however, the symbionts from *Ac. echinatior* and *T. septentrionalis* encode very similar sets of functional genes.

### Phenotypic Tests

We used our *M. whartonense S. attinicolia* cultures, and *M. lactucae* and *S. platyhelix* as controls, for several phenotypic tests. Based on the predicted gene functions, we tested for acid production from glucose, fructose, *N*-acetylglucosamine, and chitin, and for alkalization from arginine and urea. All *Mesoplasma* and *Spiroplasma* isolates produced acid from glucose and all, except *M. lactucae*, reduced the culture pH in the presence of arginine, confirming our predictions (Fig. 6). All *Mesoplasma* also produced acid from *N*-acetylglucosamine. However, no culture changed their pH in the presence of fructose, chitin, or urea.

**Figure 6).**
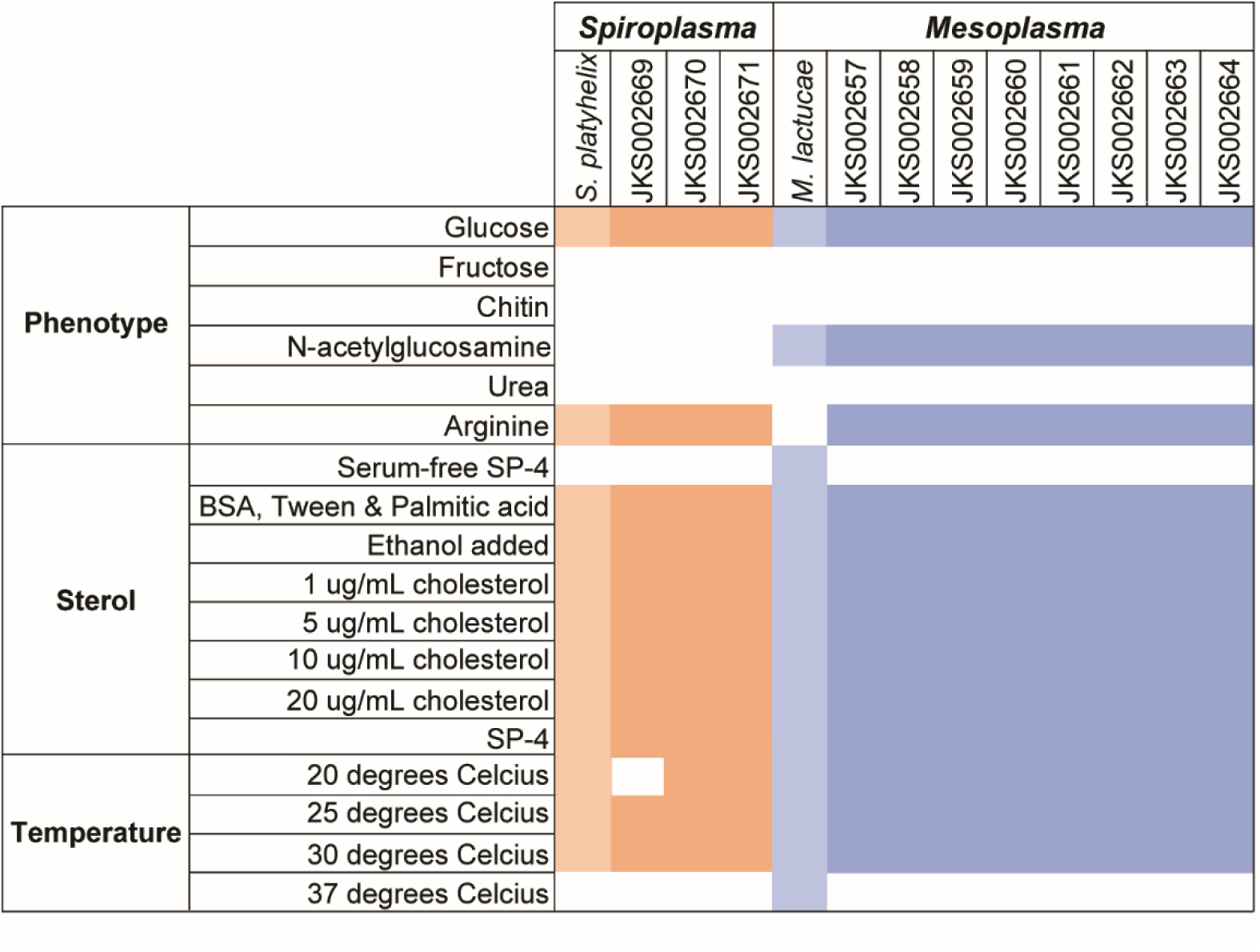
Results of phenotypic tests and growth at varying temperatures. Orange colors indicate genes present for *Spiroplasma* and blue indicates genes present for *Mesoplasma*. The lighter of each color denotes genes present for the control strain, *S. platyhelix* or *M. lactucae*. White indicates no presence.

We also tested the sterol and temperature requirements of *M. whartonense* and *S. attinicola* (Fig. 6). All of our *Mesoplasma* and *Spiroplasma* isolates grew in serum regardless of the presence of added cholesterol, but not without serum. No strains grew at 37 ℃. All *M. whartonense* isolates and *S. attinicola* strains JKS002670 and JKS002671 grew at 20, 25 and 30℃, while *Spiroplasma* strain JKS002669 only grew at 25℃ and 30℃.

Using scanning transmission electron microscopy, *M. whartonense* cells were coccoid and 150 to 200 nm in diameter (Fig. 7A). *S. attinicola* cells were non-helical rods 4 to 5 µm long (Fig. 7B).

**Figure 7).**
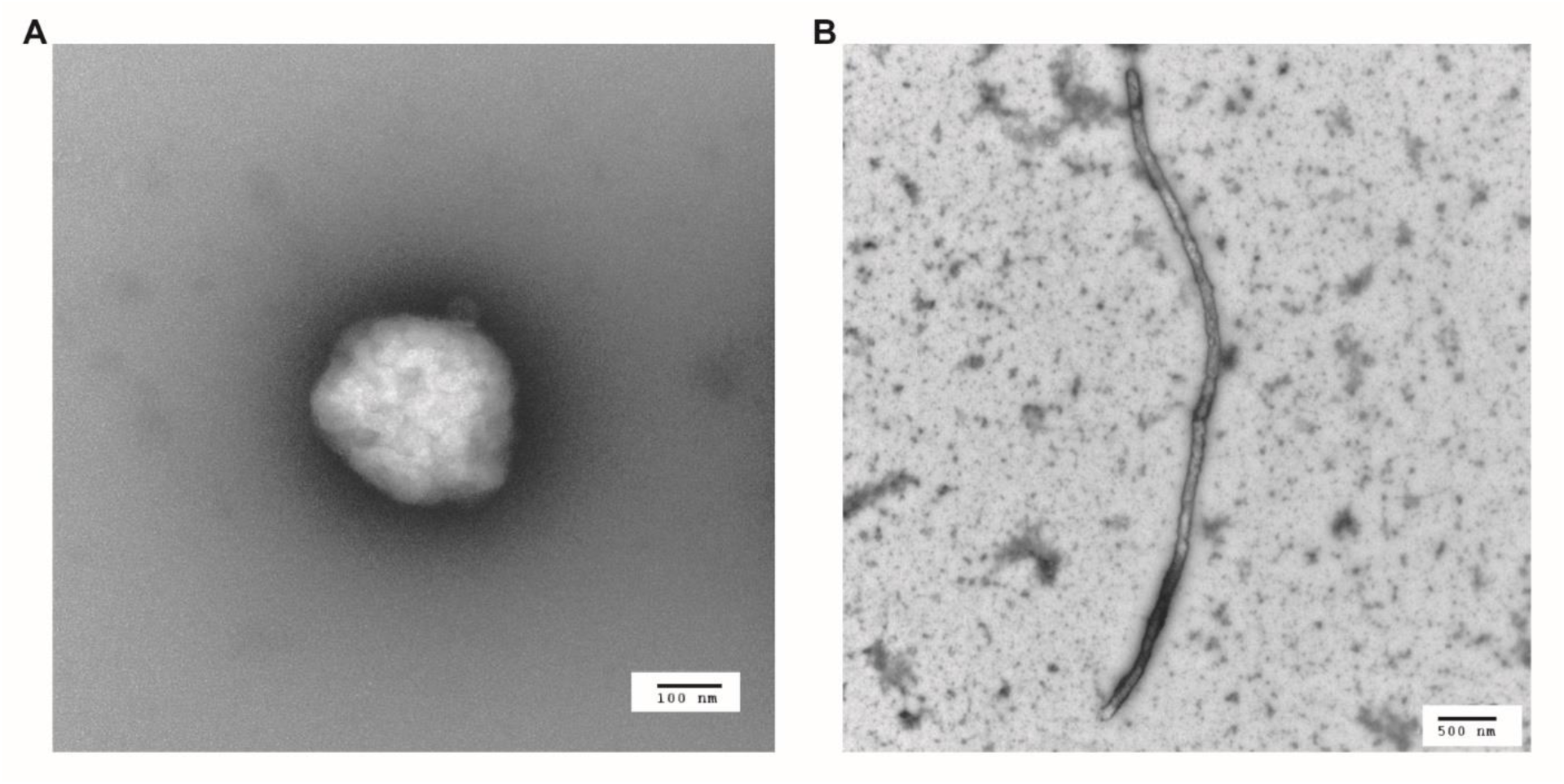
Transmission electron microscope imaging of *M. whartonense* str. JKS002660 (A) and *S. attinicola* str. JKS002670 (B)

## DISCUSSION

Here, we isolated novel *Mesoplasma* and *Spiroplasma* symbionts from the fungus-growing ant *T. septentrionalis*. They are phylogenetically related to, but distinct from, those of all previously described species and the EntAcro MAGs sequenced from *Ac. echinatior*, prompting us to describe them as *Mesoplasma whartonense sp. nov.* and *Spiroplasma attinicola sp. nov.*, respectively. Both *M. whartonense* and *S. attinicola* are predicted to encode for phenotypes that are similar to those encoded by the EntAcro MAGs, although with distinct genome sizes and genomic compositions (Fig. 4). The *M. whartonense* genomes and MAGs also phylogenetically cluster by the geographic region from which their host *T. septentrionalis* ants were collected (Fig. 2A). This contrasts to the lack of such clustering by the *T. septentrionalis Pseudonocardia* symbiont [70]. Strain specificity in the gene content of these local symbiont populations may be conserved due to adaptation to local ecological conditions such as climate and forage availability [71].

It is possible that gene content and genome size differed between the *M. whartonense* genomes and the EntAcro1 MAG may be because the EntAcro1 MAG contained extra sequences due to computational errors that occurred during contig binning from the metagenome [38, 72]. However, the *M. whartonense* MAGs assembled as part of a second study [40] had similar sizes as the *M. whartonense* isolate genomes. The difference in genomes size between the EntAcro1 and *M. whartonense* MAGs may therefore be biological or due to the different pipelines used by each group to bin their metagenomic data. If we assume that the EntAcro1 MAG accurately characterizes the genome of the *Mesoplasma* symbiont of *Ac. echinatior*, the differences in gene content and phylogenetic position between it and *M. whartonense* suggest different evolutionary paths were followed by these symbionts that are unique to each fungus-growing ant species. *Ac. echinatior* ants are found in Central America, and as leaf-cutting ants represent the most evolutionary advanced form of ant agriculture with large colonies (hundreds of thousands of ants), caste differentiation, and specialization on fresh leaves and fruits. In contrast, *T. septentrionalis* ants are found throughout the Eastern USA but belong to the “higher attine” clade that branches basal to the true leaf-cutting ants. They have smaller colonies (thousands of ants), monomorphic worker castes, and different forage preferences including insect frass, oat catkins, and leaves including oak and ferns. The cultivar fungi grown by *Ac. echinatior* and *T. septentrionalis* also largely belong to different genetic lineages. [20, 23, 24, 27, 73]. Whether Mollicute symbionts are specific to the different modes of ant agriculture or different attine ant genera is an interesting hypothesis arising from our results that will require a wide sampling of ants from different genera and geographic regions to test. Regardless, the relatively close phylogenetic relationships (belonging to the same genus at least) between either *M. whartonense* and EntAcro1 or *S. attinicola* and EntAcro10 are inconsistent with either pair co-diverging with their *T. septentrionalis* and *Ac. echinatior* ant hosts, which belong to different genera that diverged ∼19 million years ago [74].

The closest known species to *M. whartonense* and *S. attinicola* are *M. lactucae* and *S. platyhelix*, respectively, but these strains form separate groups in the phylogenetic trees (Fig. 2A & B, Suppl. Figs. S4-9) and have different gene contents (Fig. 4). *M. whartonense* and EntAcro1 both have genes for the arginine deiminase pathway, pyruvate metabolism and pyrimidine conversion, all of which *M. lactucae* lacks. The last common ancestor of *M. lactucae* and *M. whartonense*/EntAcro1 may have contained these genes, with *M. lactucae* losing them because they were not required for its symbiosis with plants. Alternatively, *M. whartonense* and EntAcro1 may have uniquely gained these genes and that were not present in their last common ancestor with *M. lactucae*. Either of these explanations would imply that *M. whartonense* and EntAcro may have retained or gained these genes to survive within the ant gut.

The *M. whartonense* genomes are also ∼100 kbps smaller than those of *M. lactucae* and MAG EntAcro1, which could mean that the many hypothetical genes in *M. lactucae* and MAG EntAcro1 were lost in *M. whartonense*. These different genome sizes could be due to *M. whartonense* losing almost all of the hypothetical genes present in MAG EntAcro1 because they are not required for the *T. septentrionalis* symbiosis, or alternatively, that these hypothetical genes were uniquely gained by MAG EntAcro1. These gene content differences might be the result of specialization of these *Mesoplasma* to each ant host, or due to technical artifacts introduced during MAG construction.

*S. attinicola* and EntAcro10 ant symbionts differ in gene content from their nearest classified neighbor, *S. platyhelix.* Both *S. attinicola* and EntAcro10 have genes for the arginine deiminase pathway, but *S. platyhelix* does not. The *S. platyhelix* genome is smaller than those of *S. attinicola* and MAG EntAcro10 but contains chitin degradation and F0F1-type ATPase genes, unlike the *Spiroplasma* ant symbionts. The last common ancestor of *S. platyhelix* and *S. attinicola*/EntAcro10 may have contained these genes, which were then lost by *S. attinicola* and EntAcro10. Alternatively, these genes may have been gained uniquely by *S. platyhelix* following its divergence from *S. attinicola* and EntAcro10. The lack of chitin degradation genes in *S. attinicola* and EntAcro10 suggest that these strains cannot digest chitin derived from the cell wall of the ants’ cultivar food source. However, this lack may benefit the ant if this chitinase could instead degrade chitin in ant tissues [75]. Differences in the distribution of hypothetical genes between the MAG EntAcro10 and *S. attinicola* could arise following the same mechanisms described above for *M. whartonense* and EntAcro1, with genes being uniquely gained or lost in either lineage. Such difference in the content of hypothetical genes between *M. whartonense* and EntAcro1 or *S. attinicola* and EntAcro10 could indicate different evolutionary adaptations to each ant host and type of fungus-growing agriculture.

*M. whartonense* and *S. attinicola* symbionts of fungus-growing ants have overlapping but distinct functions, most of which are also conserved in EntAcro1 and EntAcro10, respectively. Our gene annotations (Figs. 4 & 5) suggest that both *M. whartonense* and *S. attinicola* likely ferment glucose and fructose to produce L-lactate (both species), D-lactate and acetate (*S. attinicola*), or acetoin (*M. whartonense*). These fermentation products would be available for uptake in the ant gut or secretion onto the fungus garden. However, our phenotypic tests showed that both *M. whartonense* and *S. attinicola* strains catabolized glucose but not fructose under laboratory conditions. Only *M. whartonense* genomes possessed the genes needed degrade chitin, which might also produce acetate and ammonia that could become available to the ants or fungus garden. Under our experimental conditions *M. whartonense* degraded *N-*acetylglucosamine but not chitin.

All *M. whartonense* genomes and MAGs contained genes for citrate degradation (encoding citrate lyase; Fig. 4), as did one *S. attinicola* genome. Sapountzis et al. [39] reported citrate degradation genes in only the MAG EntAcro1 and hypothesized that citrate originated from ant foraging materials and that the acetate resulting from its degradation could be provided to the ants. Thus, both *M. whartonense* and *S. attinicola* can both likely produce acetate but from different precursors (citrate and pyruvate). However, how citrate degradation benefits either *M. whartonense* and *S. attinicola* remains vague, because genes are lacking that might encode further transformations of oxaloacetate that might provide them an energetic benefit. Biochemical evidence for malate dehydrogenase activity (which could regenerate NAD^+^ by reducing oxaloacetate) exists for many mollicutes, despite their lacking homologs to known malate dehydrogenase genes [69]. The benefit of citrate metabolism for *M. whartonense* (and the one *S. attinicola* strain) and the degree to which it provides acetate for the ant host therefore remains unclear.

*S. attinicola*, but not *M. whartonense* encodes genes for glycerol degradation (Fig. 4), which is tied to pathogenicity in *Mycoplasma*, another Mollicute genus, via the production of hydrogen peroxide [76]. However, *M. whartonense* and *S. attinicola* lack genes coding for GlpO, which is needed to produce hydrogen peroxide. Glycerol likely originates from the ant host and can be broken down into glyceraldehyde-3-phosphate that enters the glycolysis or pentose phosphate pathways. Glycerol may be produced by the lipases encoded by *M. whartonense* and *S. attinicola* (by which they presumably obtain host lipids for membrane biogenesis) and/or used for glycerophospholipid biosynthesis by *M. whartonense* (but not *S. attinicola*, which lacks the corresponding genes).

Both *M. whartonense* and *S. attinicola* strains all possess the genes needed to degrade arginine (via the arginine deiminase pathway; Fig. 4) and could do so in culture (Fig. 6). Although present in many Mollicutes, these genes are present in only one other *Mesoplasma* genome (*M. photuris*) but are sometimes found in *Spiroplasma* genomes [77]. Both *Ac. echinatior* and *T. septentrionalis* ants lack genes for arginine metabolism and receive this amino acid from their fungal cultivar [78, 79]. The degradation of arginine produces ammonium, and Sapountzis et al [39] hypothesized that EntAcro1 and EntAcro10 recycled excess arginine into ammonium and provided it to the fungus garden for use in protein synthesis. This hypothesis was supported by these bacteria expressing arginine transporters in the ant hindgut and the presence of ammonium in *Ac. echinatior* ant fecal droplets [39]. Our study also supports this hypothesis by showing that *M. whartonense* and *S. attinicola* catabolize arginine in culture. However, glycolytic Mollicutes (such as *M. whartonense* and *S. attinicola*) can generate ATP via glucose hydrolysis, and therefore may not require the high flux through the arginine deiminase pathway to generate ATP needed by non-glycolytic Mollicute species [69], concomitantly producing less ammonium in ant guts. The arginine deiminase pathway can also help Mollicutes overcome acid stress [80], consistent with the low pH values measured for ant guts [81]. Determining the function of the arginine deiminase pathway in ant guts and if its products are, in fact, provided to the fungus garden will require future experiments.

Our isolation of the novel ant symbionts *M. whartonense* and *S. attinicola* sets the stage for future research removing and reintroducing both Mollicutes to the ants individually and in competition will determine why these two bacterial species do not co-occur in the same ant [35]. They will also facilitate other co-culture assays test if these species interact with other ant or fungus garden symbionts. Most critically, they will facilitate tests of what role *M. whartonense* and *S. attinicola* play in the fungus-growing ant symbiosis. Sapountzis et al. [39] previously proposed that EntAcro1 and EntAcro10 benefited the symbiosis by producing acetate and ammonium that could be transferred to the ant or fungus garden. Although we agree with these potential benefits, future work will determine if ecologically significant amounts of either product are made *in situ* and if their production offsets the loss of other metabolites that *M. whartonense* and *S. attinicola* acquire from the ants or ant food (e.g., lipids, nucleobases, vitamins, arginine [primarily for catabolism] and other amino acids [for anabolism], and other carbon sources). It remains possible that the cost of consuming these nutrients exceeds the provision of benefits to the ant host, more consistent with parasitism than mutualism, or that non-nutritive benefits are involved, e.g., colonization resistance or immune priming.

Finally, we found that the predicted functions encoded by *M. whartonense* and *S. attinicola* very similar to those encoded by MAGs EntAcro1 and EntAcro10, respectively, although the *M. whartonense* genomes are smaller than MAG EntAcro1 and all genomes from *T. septentrionalis* ants are distinct from the MAGs from *Ac. echinatior*. The future characterization of *M. whartonense-* and *S. attinicola*-like symbionts from *Ac. echinatior* and other Attine ants will reveal their host specificity, coevolution, and regional differentiation. In doing so, they will reveal the full diversity and function of these enigmatic members of the ant-microbe symbiosis.

### Description of Mesoplasma whartonense

Etymology: whartonense, neutr. of Wharton State Park, referring to the geographic location of where the ant was collected from which the bacterium was isolated.

Isolated from the *T. septentrionalis* gut in Wharton State Park New Jersey USA, with no known pathogenicity. Cells are coccoid in shape, 150 to 200 nm in size, lack a cell wall, and are filterable through 0.45 µm membranes. Colonies are a fried-egg shape, grow on SP-4 agar in 4 to 7 days at 30 °C, and can grow between 20 and 30°C. Produces acid from glucose and *N*-acetylglucosamine, and hydrolyzes arginine but not urea. Requires serum for growth but not cholesterol. The 16S rRNA gene of the proposed type strain (JKS002660) was 93.45% similar to that of *Mesoplasma lactucae* 831-C4^T^. The genome of *Mesoplasma whartonense* JKS002660 has 72.80% Average Nucleotide Identity to that of *M. lactucae* 831-C4^T^. Its genome size is 786,541 bp and its %GC content is 32.68%. The type strain is JKS002660 (NCTC 14863; DMSZ 115716) and other strains include JKS002657, JKS002658, JKS002659, JKS002661, JKS002662, JKS002663, and JKS002664.

### Description of Spiroplasma attinicola

Etymology: attini- Attine, -cola neutr. Inhabitant/dweller. An inhabitant of a fungus-growing ant from Tribe Attini

Isolated from the *T. septentrionalis* gut in Wharton State Park New Jersey USA, with no known pathogenicity. Cells are rod and non-helical in shape, 4 to 5 µm long, lack a cell wall, and are filterable through 0.45 µm membranes. Colonies are a fried-egg shape, grow on SP-4 agar in 7 days at 30 °C, and can grow between 20 and 30 °C. Produces acid from glucose and does not hydrolyze arginine or urea. Requires serum for growth but not cholesterol. The 16S rRNA gene of the proposed type strain (JKS002670) was 94.12% similar to *Spiroplasma platyhelix*. The genome of *Spiroplasma attinicola* has 77.63% Average Nucleotide Identity to *S. platyhelix*. Its genome size is 872,740 bp and its %GC content is 25.49%. The type strain is JKS002670 (NCTC 14964; DMSZ 115717) and other strains include JKS002669 and JKS002671.

## Acknowledgements

We thank the New Jersey State Forest staff for their assistance with our fieldwork, and the members of the Klassen lab for help collecting colonies and thoughtful comments on the paper. We also thank Dr. Kendra Maas and University of Connecticut Microbiology Analysis, Resources, and Services facility for sequencing and Dr. Xuanhao Sun at University of Connecticut’s Bioscience Electron Microscopy Laboratory for imaging. This research was funded by the National Science Foundation (IOS-1656475) and the University of Connecticut.

## Notes

### Competing Interest Statement

The authors have declared no competing interest.

