## Supplementary Figures and Tables for "Isolation and characterization of mollicute symbionts from a fungus-growing ant reveals genome reduction and host specialization"

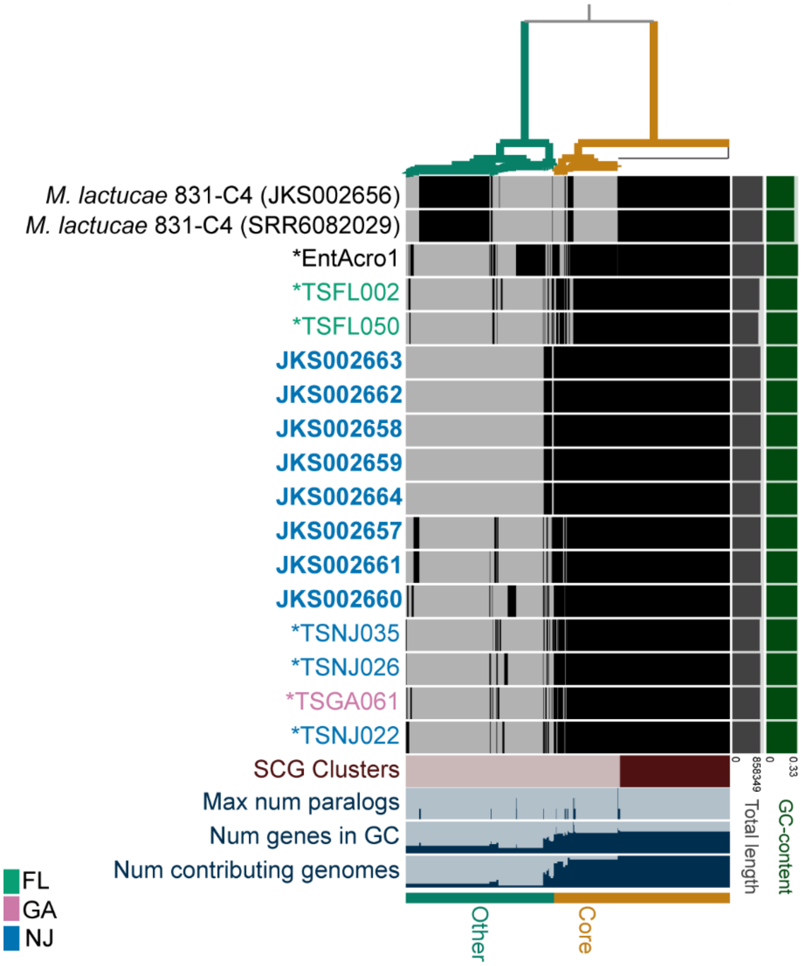


Supplementary Figure S1) Anvi'o pangenome analysis of *M. whartonense* genomes (bold) and MAGs with a 100% BUSCO score (*), the EntoAcro1 MAG (*), and the *M. lactucae* genomes as a reference. Text colors indicate strains or MAGs from the same state. Two *M. lactucae* genomes are shown, representing the genome downloaded from NCBI (SRR6082029) and the genome that we re-sequenced as a control (JKS002656). In the central heatmap, black and grey bars indicate homologous genes that are present or absent in each genome. The dendrogram clusters groups of homologous genes (anvi'o gene clusters) using Euclidean distances.

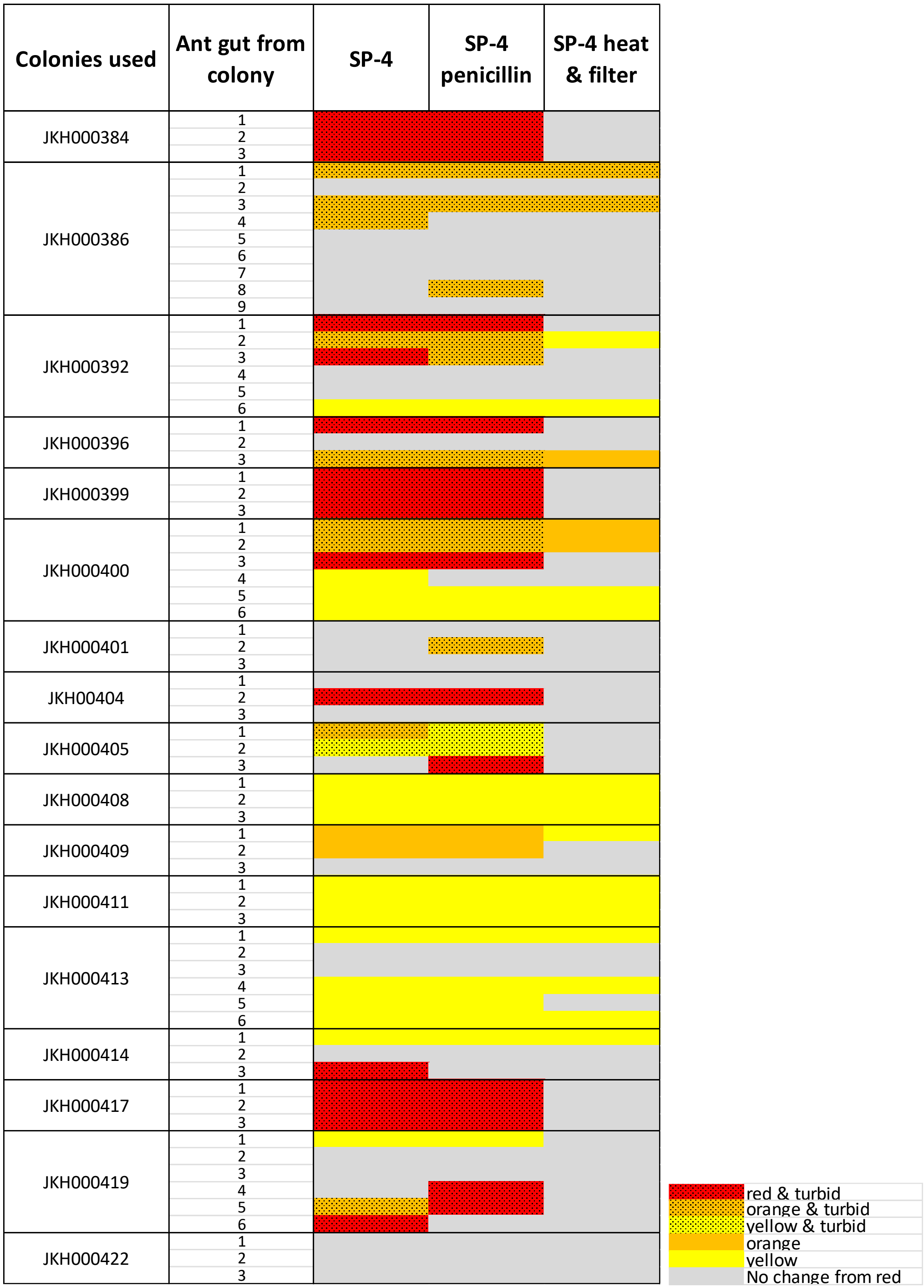


Supplementary Figure S2) Heatmap describing the cultures used to enrich *Mesoplasma* and *Spiroplasma* from ant guts.

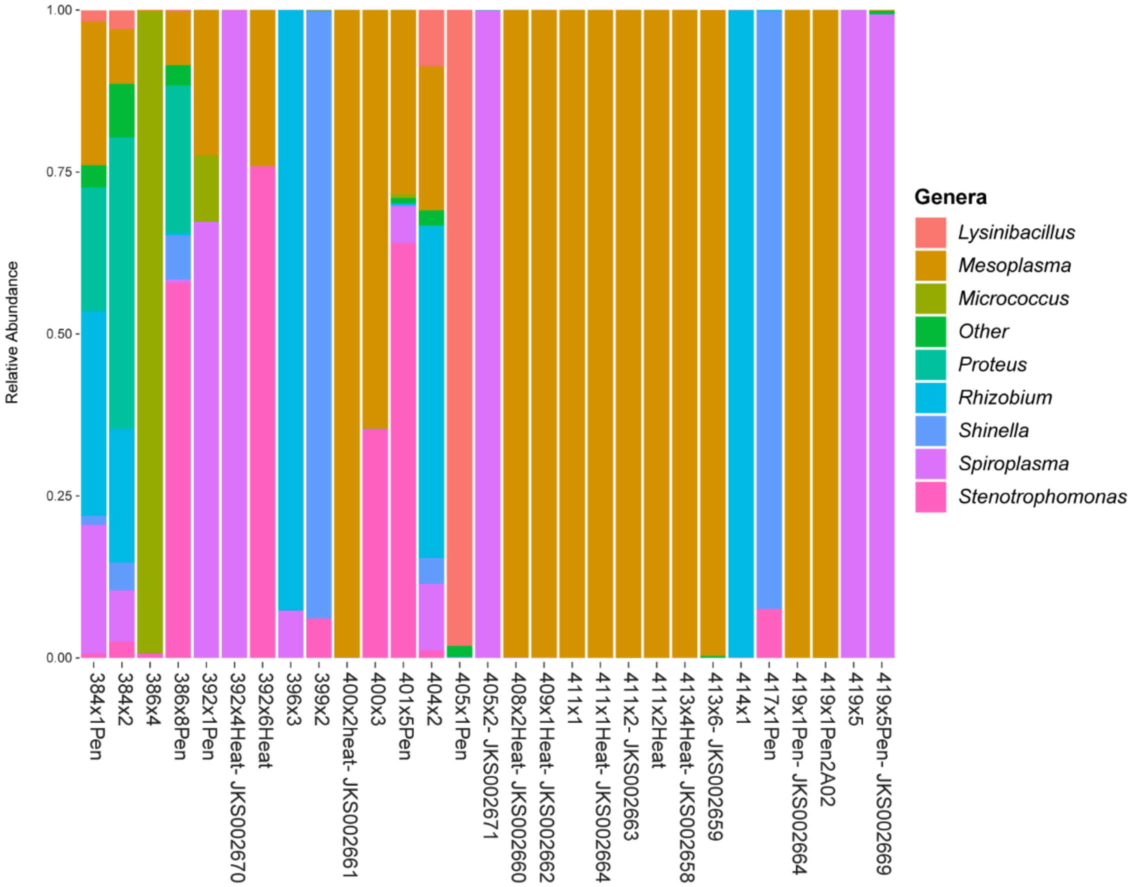


Supplementary Figure S3) Bar plot representing the top genera present ( ≥ 15% abundance) in our 16S rRNA community amplicon sequencing of cultures. Each bar represents an individual culture. On the x-axis, names with a JKS prefix indicate samples that were used isolation and genome sequencing.

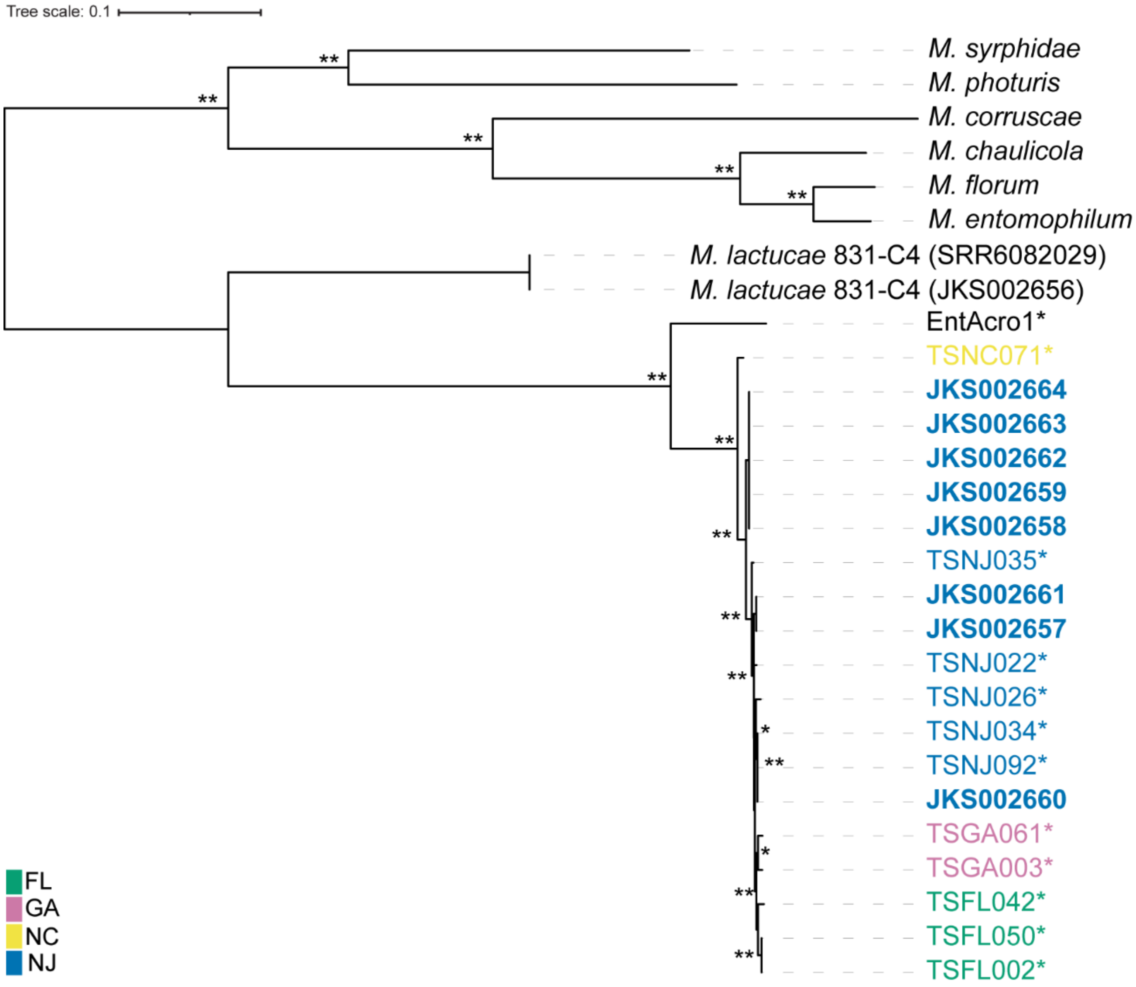


Supplementary Figure S4) *Mesoplasma* amino acid phylogeny created using 21 single-copy core gene clusters as input data and the fasttree WAG + CAT substitution model. Asterisks indicate MAGs and bolded names indicate *M. whartonense* isolate genomes. Colors group strains and MAGs by state. The tree was rooted using the midpoint of the branch between *M. lactucae* and the other reference genomes. Local bootstrap values of 60–79 and 80–100 are indicated by * and **, respectively.

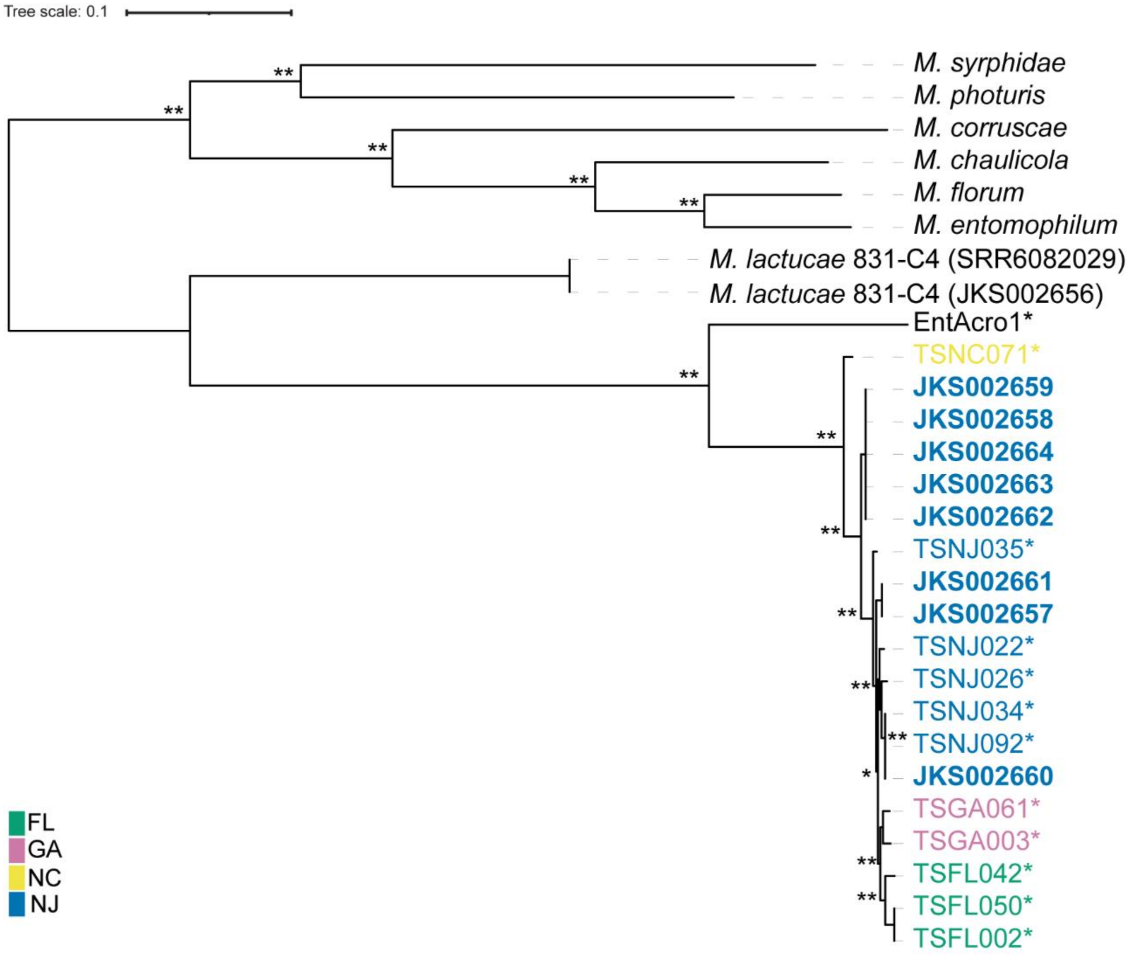


Supplementary Figure S5) *Mesoplasma* nucleotide phylogeny created using 21 single-copy core gene clusters as input data and the fasttree JC + CAT substitution model. Asterisks indicate MAGs and bolded names indicate *M. whartonense* isolate genomes. Colors group strains and MAGs by state. The tree was rooted using the midpoint of the branch between *M. lactucae* and the other reference genomes. Local bootstrap values of 60–79 and 80–100 are indicated by * and **, respectively.

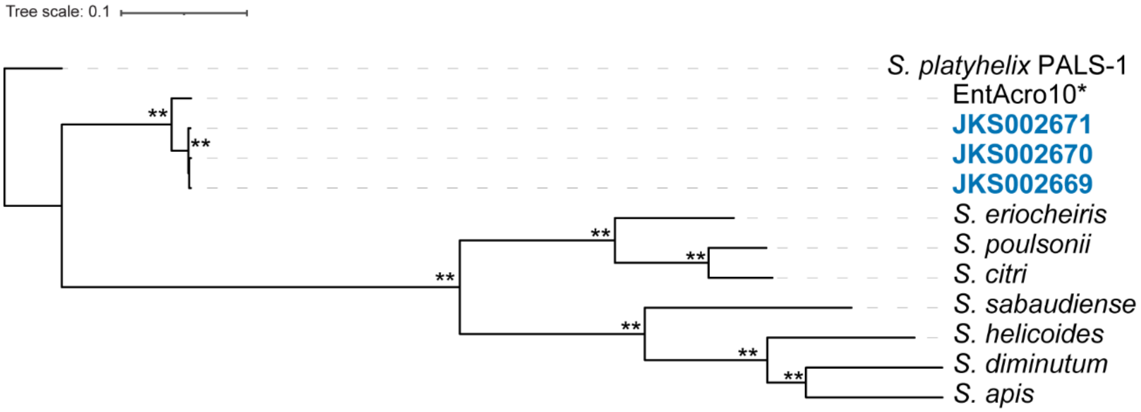


Supplementary Figure S6) *Spiroplasma* amino acid phylogeny created using 100 single-copy core gene clusters as input data and the fasttree WAG + CAT substitution model. Asterisks indicate MAGs and bolded names indicate *S. attinicola* isolate genomes. The blue color indicates strains from New Jersey. The tree was rooted using the *S. platyhelix* genome. Local bootstrap values of 60–79 and 80–100 are indicated by * and **, respectively.

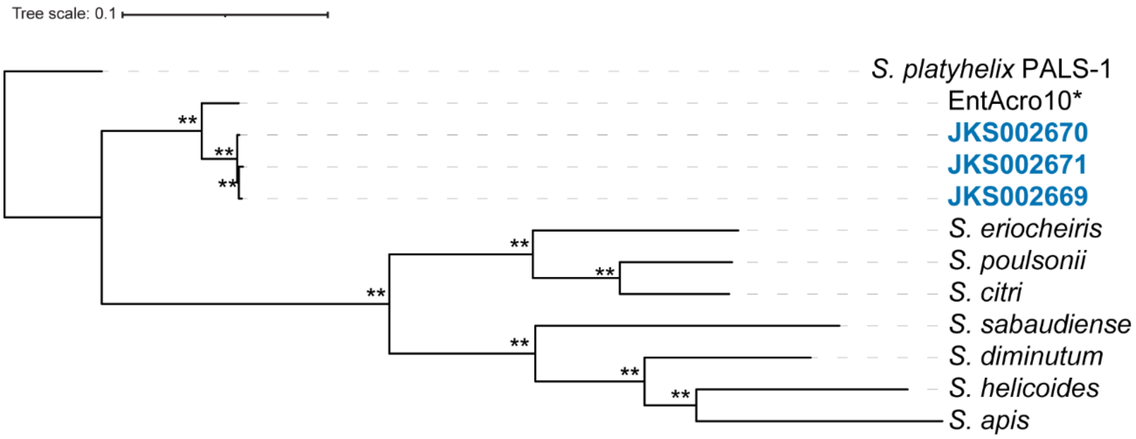


Supplementary Figure S7) *Spiroplasma* nucleotide phylogeny created using 100 single-copy core gene clusters as input data and the fasttree JC + CAT substitution model. Asterisks indicate MAGs and bolded names indicate *S. attinicola* isolate genomes. The blue color indicates strains from New Jersey. The tree was rooted using the *S. platyhelix* genomes. Local bootstrap values of 60–79 and 80–100 are indicated by * and **, respectively.

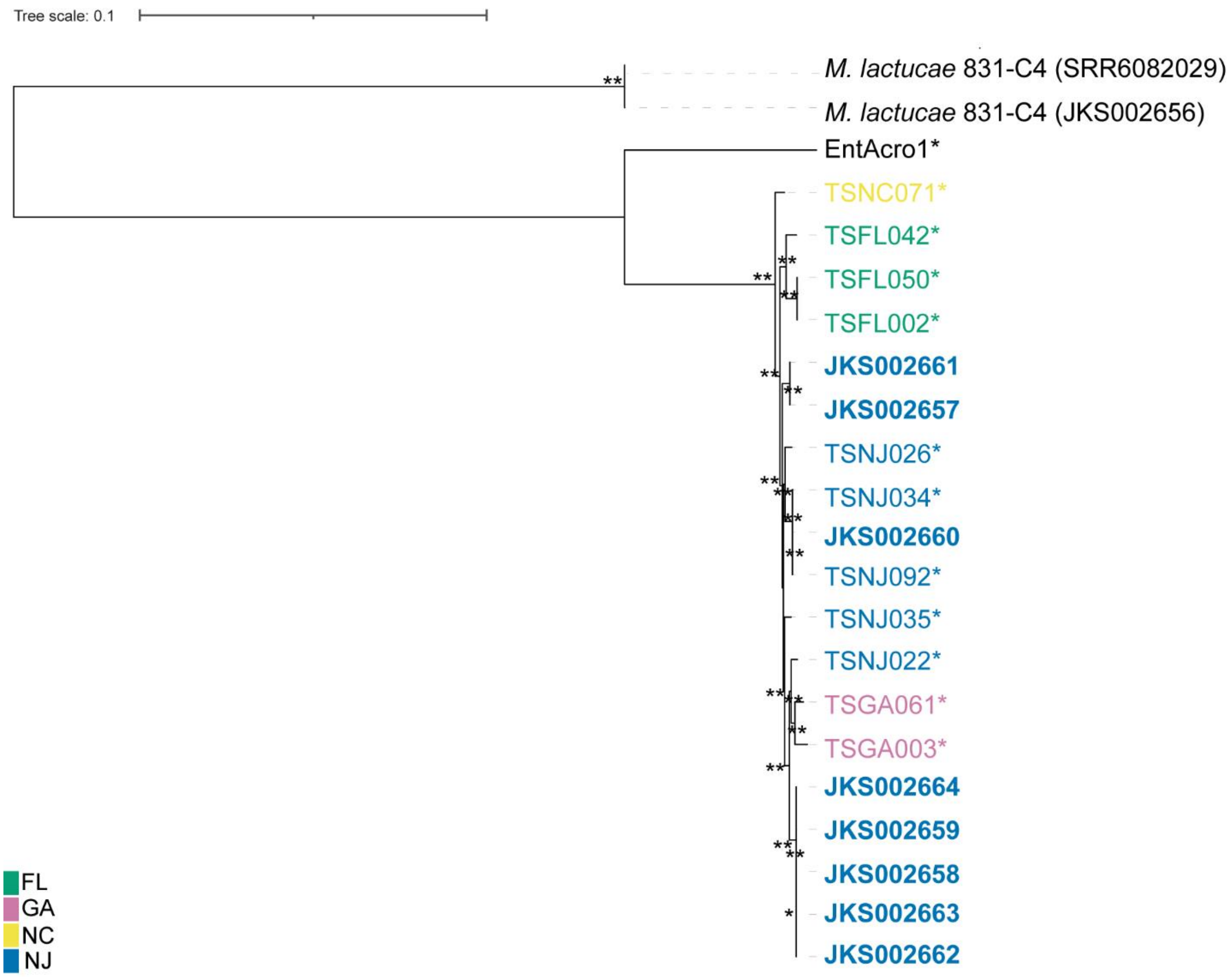


Supplementary Figure S8) *Mesoplasma* amino acid phylogeny created using 550 core gene clusters as input data and the fasttree WAG + CAT substitution model. Asterisks indicate MAGs and bolded names indicate *M. whartonense* isolate genomes. Colors group strains and MAGs by state. The tree was rooted using the midpoint of the branch between *M. lactucae* genomes. Local bootstrap values of 60–79 and 80–100 are indicated by * and **, respectively.

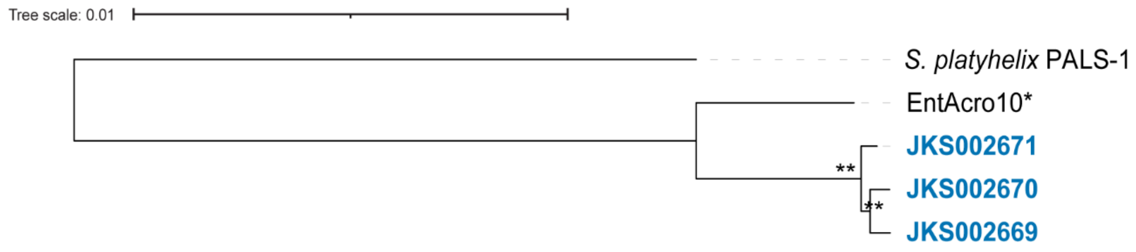

Supplementary Figure S9) *Spiroplasma* amino acid phylogeny created using 684 core gene clusters as input data and the fastree WAG + CAT substitution model. Asterisks indicate MAGs and bolded names indicate *S. attinicola* isolate genomes, with the blue text indicating strains isolated from New Jersey. The tree was rooted using the *S. platyhelix* genome. Local bootstrap values of 60–79 and 80–100 are indicated by * and **, respectively.

**Supplemental Tables**

Supplementary Table S1) Colony ID, date and location with the respective Genome or MAG ID associated with that colony.

| Colony ID | Collection Date | Geographic Location | GPS Latitude | GPS Longitude | Genome or MAG ID |
|---|---|---|---|---|---|
| JKH000014 | 5/20/2013 | Florida:Wekiwa Springs State Park | 28.70991 | -81.46795 | TSFL002 |
| JKH000065 | 7/22/2014 | New Jersey:Brendon T. Byrne State Forest | 39.87275 | -74.52157 | TSNJ022 |
| JKH000069 | 7/22/2014 | New Jersey:Brendon T. Byrne State Forest | 39.87212 | -74.52075 | TSNJ026 |
| JKH000073 | 7/23/2014 | New Jersey:Wharton State Forest | 39.71267 | -74.5635 | TSNJ092 |
| JKH000079 | 7/23/2014 | New Jersey:Wharton State Forest | 39.71275 | -74.56442 | TSNJ034 |
| JKH000082 | 7/24/2014 | New Jersey:Wharton State Forest:Goshen pond | 39.74587 | -74.7615 | TSNJ035 |
| JKH000105 | 11/14/2014 | Florida:Wekiwa Springs | 28.709 | -81.4873 | TSFL042 |
| JKH000119 | 11/16/2014 | Florida:Paynes | 27.62286 | -81.8037 | TSFL050 |
| JKH000136 | 5/7/2015 | Geogia:Yuchi Wildlife Management Area | 33.08388 | -81.77029 | TSGA003 |
| JKH000138 | 5/7/2015 | Geogia:Yuchi Wildlife Management Area | 33.08393 | -81.76996 | TSGA061 |
| JKH000154 | 6/9/2015 | North Carolina:William B Umpstead State Park | 35.8607 | -78.76103 | TSNC071 |
| JKH000392 | 6/23/2020 | New Jersey:Wharton State Forest:Quaker Bridge | 39.70921 | -74.66321 | JKS002670 |
| JKH000400 | 6/24/2020 | New Jersey:Wharton State Forest:Batona Field | 39.78143 | -74.62846 | JKS002661 |
| JKH000405 | 6/24/2020 | New Jersey:Wharton State Forest:Batona Field | 39.78159 | -74.62822 | JKS002671 |
| JKH000408 | 6/25/2020 | New Jersey:Wharton State Forest:Hawkins Bridge | 39.71257 | -74.56377 | JKS002660 |
| JKH000409 | 6/25/2020 | New Jersey:Wharton State Forest:Hawkins Bridge | 39.71259 | -74.56346 | JKS002662 |
| JKH000411 | 6/25/2020 | New Jersey:Wharton State Forest:Hawkins Bridge | 39.71284 | -74.56367 | JKS002663 & JKS002664 |
| JKH000413 | 6/25/2020 | New Jersey:Wharton State Forest:Hawkins Bridge | 39.71269 | -74.56355 | JKS002658 & JKS002659 |
| JKH000419 | 6/26/2020 | New Jersey:Brendan Byrne State Forest | 39.87226 | -74.52554 | JKS002657 & JKS002669 |

Supplementary Table S2) Characteristics of the *T. septentrionalis Mesoplasma* MAGs sequenced by JGI.

| Name | Colony ID | Length (bp) | BUSCO completeness score | Contigs | % GC | JGI Bin ID |
|---|---|---|---|---|---|---|
| TSFL002 | JKH000014 | 738626 | 94.70% | 45 | 32.72% | 3300012079_26 |
| TSFL042 | JKH000105 | 615878 | 67.30% | 36 | 32.41% | 3300012141_10 |
| TSFL050 | JKH000119 | 738262 | 94.70% | 45 | 32.72% | 3300012098_5 |
| TSGA003 | JKH000136 | 524149 | 72% | 20 | 32.56% | 3300012101_6 |
| TSGA061 | JKH000138 | 752725 | 100% | 9 | 32.62% | 3300012067_9 |
| TSNC071 | JKH000154 | 642949 | 82% | 8 | 32.58% | 3300011392_7 |
| TSNJ022 | JKH000065 | 758990 | 100% | 10 | 32.51% | 3300012151_17 |
| TSNJ026 | JKH000069 | 760750 | 100% | 9 | 32.55% | 3300012165_16 |
| TSNJ034 | JKH000079 | 385824 | 52.70% | 77 | 33% | 3300012155_29 |
| TSNJ035 | JKH000082 | 744761 | 100% | 9 | 32.57% | 3300011404_8 |
| TSNJ092 | JKH000073 | 612061 | 69.30% | 9 | 32.55% | 3300012100_6 |

Supplementary Table S3) Characteristics of the *Mesoplasma* and *Spiroplasma* MAGs published by Sapountzis et al. [39]

| Name | Length (bp) | BUSCO completeness score | Contigs | % GC |
|---|---|---|---|---|
| EntAcro1 | 866917 | 100% | 16 | 33.74% |
| EntAcro10 | 838545 | 98% | 13 | 25.53% |

Supplementary Table S4) Characteristics of the *Mesoplasma* and *Spiroplasma* reference genomes downloaded from NCBI.

| **Name** | **Strain** | **ATCC ID #** | **Assembly** | **BioSample** | **BioProject** | **Size (bp)** |
|---|---|---|---|---|---|---:|
| *Mesoplasma florum* | L1 | ATCC:33453 | GCF_000008305.1 | SAMN03081416 | PRJNA224116 | 793,224 |
| *Mesoplasma corruscae* | ELCA-2 | | GCA_002930145.1 | SAMN05436728 | PRJNA331058 | 839,085 |
| *Mesoplasma chauliocola* | CHPA-2 | | GCF_002290085.1 | SAMN07573570 | PRJNA224116 | 854,780 |
| *Mesoplasma photuris* | | ATCC:49581 | GCA_000702725.1 | SAMN02841188 | PRJNA223039 | 778,966 |
| *Mesoplasma syrphidae* | YJS | ATCC:51578 | GCF_002843565.1 | SAMN08158127 | PRJNA224116 | 908,214 |
| *Mesoplasma entomophilum* | TAC | | GCF_002749675.1 | SAMN07838951 | PRJNA224116 | 847,967 |
| *Mesoplasma lactucae* | 831-C4 | ATCC:49193 | GCA_002441935.1 | SAMN07709945 | PRJNA412357 | 837,471 |
| *Spiroplasma citri* | R8-A2 | | GCF_001886855.1 | SAMN04110376 | PRJNA296877 | 1,599,709 |
| *Spiroplasma eriocheiris* | DSM 21848 | | GCA_001029265.1 | SAMN03488050 | PRJNA253647 | 1,365,714 |
| *Spiroplasma sabaudiense* | Ar-1343 | | GCA_000565215.1 | SAMN03081504 | PRJNA184750 | 1,075,953 |
| *Spiroplasma poulsonii* | MSRO_BK | | GCF_009866525.1 | SAMN11349289 | PRJNA224116 | 1,938,611 |
| *Spiroplasma apis* | B31 | ATCC:33834 | GCA_000500935.1 | SAMN02641479 | PRJNA184751 | 1160554 |
| *Spiroplasma diminutum* | CUAS-1 | | GCA_000439455.1 | SAMN02602967 | PRJNA184745 | 945296 |
| *Spiroplasma helicoides* | TABS-2 | | GCA_001715535.1 | SAMN05578878 | PRJNA253649 | 1326546 |
| *Spiroplasma platyhelix* | PALS-1 | ATCC 51748 | GCA_012163225.1 | SAMN14173566 | PRJNA608455 | 739,863 |

Supplementary Table S5) Characteristics of the *M. whartonense* and *S. attinicola* isolate genomes and *M. lactucae* reference genomes sequenced in this study.

| | Strain ID | Sample ID | Colony ID | Contigs before filtering | Contigs after filtering | % GC | BUSCO completeness score[a] | Largest contig (bp) | Total length (bp) | NCBI WGS accession |
|---|---|---|---|---|---|---|---|---|---|---|
| *Mesoplasma* | JKS002656 | | | 8 | 8 | 29.63 | 99.30% | 443,977 | 824,849 | JAKNSY000000000 |
| | JKS002657 | | JKH000419 | 8 | 5 | 32.74 | 100% | 308,895 | 769,567 | JAKNSX000000000 |
| | JKS002658 | JKA007839 | JKH000413 | 12 | 7 | 32.79 | 100% | 448,687 | 767,909 | JAKNSW000000000 |
| | JKS002659 | JKA007841 | JKH000413 | 10 | 6 | 32.79 | 100% | 448,687 | 770,321 | JAKNSV000000000 |
| | JKS002660 | JKA007836 | JKH000408 | 8 | 6 | 32.68 | 100% | 545,137 | 786,541 | JAKNSU000000000 |
| | JKS002661 | JKA007851 | JKH000400 | 6 | 5 | 32.74 | 100% | 284,762 | 769,577 | JAKNST000000000 |
| | JKS002662 | JKA007834 | JKH000409 | 12 | 7 | 32.79 | 100% | 448,689 | 767,977 | JAKNSS000000000 |
| | JKS002663 | JKA007853 | JKH000411 | 14 | 9 | 32.70 | 100% | 221,966 | 762,874 | JAKNSR000000000 |
| | JKS002664 | JKA007852 | JKH000411 | 13 | 9 | 32.72 | 100% | 221,966 | 765,176 | JAKNSQ000000000 |
| *Spiroplasma* | JKS002669 | JKA007833 | JKH000419 | 12 | 4 | 25.7 | 98.70% | 632,388 | 835,789 | JAKQXY000000000 |
| | JKS002670 | JKA007855 | JKH000392 | 18 | 8 | 25.49 | 98.70% | 372,052 | 872,740 | JAKNSP000000000 |
| | JKS002671 | JKA007849 | JKH000405 | 10 | 5 | 25.77 | 98.70% | 528,435 | 884,092 | JAKNSO000000000 |

[a]BUSCO scores were determined for the filtered contigs

Supplementary Table S6) Classic RAST Overview of the *M. whartonense* and *S. attinicola* genomes, *Mesoplasma* MAGs, EntAcro MAGs, and the *M. lactucae* and *S. platyhelix* reference genomes.

| | *Mesoplasma* | | | | | | | | | | | | | | *Spiroplasma* | | | | |
|---|---|---|---|---|---|---|---|---|---|---|---|---|---|---|---|---|---|---|---|
| **RAST Annotation** | **EnAcro1** | **JKS002657** | **JKS002658** | **JKS002659** | **JKS002660** | **JKS002661** | **JKS002662** | **JKS002663** | **JKS002664** | **TSGA061** | **TSNJ022** | **TSNJ026** | ***M. lactucae*** | **TSNJ035** | **EntAcro10** | **JKS002669** | **JKS002670** | **JKS002671** | ***S. platyhelix*** |
| Number of Subsystems | 152 | 147 | 151 | 151 | 152 | 147 | 151 | 152 | 152 | 148 | 151 | 150 | 148 | 148 | 141 | 139 | 140 | 140 | 135 |
| Number of Coding Sequences | 757 | 671 | 661 | 666 | 689 | 668 | 662 | 660 | 663 | 655 | 663 | 663 | 691 | 654 | 774 | 768 | 820 | 814 | 670 |
| Number of RNAs | 35 | 34 | 37 | 37 | 34 | 34 | 37 | 34 | 34 | 35 | 33 | 33 | 35 | 33 | 29 | 29 | 29 | 29 | 29 |
| Cofactors, Vitamins, Prosthetic Groups, Pigments | 42 | 43 | 43 | 42 | 43 | 43 | 43 | 43 | 43 | 43 | 44 | 43 | 48 | 43 | 46 | 35 | 35 | 35 | 24 |
| Cell Wall and Capsule | 4 | 4 | 4 | 4 | 4 | 4 | 4 | 4 | 4 | 4 | 4 | 4 | 4 | 4 | 2 | 2 | 2 | 2 | 2 |
| Virulence, Disease and Defense | 16 | 16 | 20 | 19 | 20 | 16 | 20 | 20 | 20 | 16 | 21 | 20 | 19 | 20 | 23 | 18 | 18 | 18 | 17 |
| Potassium metabolism | 6 | 2 | 2 | 2 | 2 | 2 | 2 | 2 | 2 | 2 | 2 | 2 | 6 | 2 | 2 | 2 | 2 | 2 | 2 |
| Photosynthesis | 0 | 0 | 0 | 0 | 0 | 0 | 0 | 0 | 0 | 0 | 0 | 0 | 0 | 0 | 0 | 0 | 0 | 0 | 0 |
| Miscellaneous | 1 | 1 | 1 | 1 | 1 | 1 | 1 | 1 | 1 | 1 | 1 | 1 | 1 | 1 | 1 | 1 | 1 | 1 | 1 |
| Phages, Prophages, Transposable elements, Plasmids | 0 | 0 | 0 | 0 | 0 | 0 | 0 | 0 | 0 | 0 | 0 | 0 | 0 | 0 | 0 | 0 | 0 | 0 | 0 |
| Membrane Transport | 19 | 20 | 20 | 20 | 20 | 20 | 20 | 20 | 20 | 20 | 20 | 20 | 22 | 20 | 13 | 14 | 12 | 15 | 15 |
| Iron acquisition and metabolism | 0 | 0 | 0 | 0 | 0 | 0 | 0 | 0 | 0 | 0 | 0 | 0 | 0 | 0 | 0 | 0 | 0 | 0 | 0 |
| RNA Metabolism | 50 | 49 | 49 | 49 | 49 | 49 | 49 | 49 | 49 | 49 | 49 | 49 | 45 | 49 | 47 | 46 | 46 | 46 | 44 |
| Nucleosides and Nucleotides | 27 | 27 | 27 | 27 | 27 | 27 | 27 | 27 | 27 | 26 | 27 | 27 | 20 | 26 | 28 | 29 | 29 | 29 | 28 |
| Protein Metabolism | 137 | 117 | 117 | 139 | 139 | 117 | 117 | 139 | 139 | 139 | 139 | 139 | 143 | 117 | 132 | 134 | 133 | 132 | 132 |
| Cell Division and Cell Cycle | 11 | 12 | 12 | 12 | 12 | 12 | 12 | 12 | 12 | 12 | 12 | 12 | 9 | 12 | 15 | 15 | 15 | 15 | 15 |
| Motility and Chemotaxis | 0 | 0 | 0 | 0 | 0 | 0 | 0 | 0 | 0 | 0 | 0 | 0 | 0 | 0 | 0 | 0 | 0 | 0 | 0 |
| Regulation and Cell signaling | 3 | 3 | 3 | 3 | 3 | 3 | 3 | 3 | 3 | 3 | 3 | 3 | 3 | 3 | 2 | 2 | 2 | 2 | 2 |
| Secondary Metabolism | 0 | 0 | 3 | 3 | 3 | 0 | 3 | 3 | 3 | 0 | 3 | 3 | 0 | 3 | 0 | 0 | 0 | 0 | 0 |
| DNA Metabolism | 45 | 39 | 50 | 48 | 48 | 39 | 50 | 50 | 50 | 42 | 44 | 41 | 40 | 36 | 38 | 42 | 42 | 44 | 35 |
| Fatty Acids, Lipids, and Isoprenoids | 12 | 12 | 12 | 12 | 12 | 12 | 12 | 12 | 12 | 12 | 12 | 12 | 14 | 12 | 3 | 3 | 3 | 3 | 5 |
| Nitrogen Metabolism | 0 | 0 | 0 | 0 | 0 | 0 | 0 | 0 | 0 | 0 | 0 | 0 | 0 | 0 | 0 | 0 | 0 | 0 | 0 |
| Dormancy and Sporulation | 1 | 1 | 1 | 1 | 1 | 1 | 1 | 1 | 1 | 1 | 1 | 1 | 1 | 1 | 1 | 1 | 1 | 1 | 1 |
| Respiration | 8 | 8 | 8 | 8 | 8 | 8 | 8 | 8 | 8 | 8 | 8 | 8 | 9 | 8 | 3 | 3 | 3 | 3 | 9 |
| Stress Response | 10 | 10 | 19 | 10 | 10 | 10 | 10 | 10 | 10 | 10 | 10 | 10 | 12 | 10 | 4 | 3 | 3 | 12 | 3 |
| Metabolism of Aromatic Compounds | 0 | 0 | 9 | 0 | 0 | 0 | 0 | 0 | 0 | 0 | 0 | 0 | 0 | 0 | 0 | 0 | 0 | 0 | 0 |
| Amino Acids and Derivatives | 21 | 22 | 24 | 24 | 23 | 22 | 24 | 24 | 24 | 23 | 23 | 23 | 6 | 23 | 20 | 20 | 20 | 20 | 17 |
| Sulfur Metabolism | 2 | 2 | 2 | 2 | 2 | 2 | 2 | 2 | 2 | 2 | 2 | 2 | 3 | 2 | 2 | 1 | 1 | 1 | 1 |
| Phosphorus Metabolism | 11 | 11 | 11 | 11 | 11 | 11 | 11 | 11 | 11 | 11 | 11 | 11 | 11 | 11 | 5 | 5 | 5 | 5 | 6 |
| Carbohydrates | 46 | 47 | 46 | 47 | 46 | 46 | 46 | 46 | 47 | 46 | 48 | 48 | 43 | 46 | 49 | 47 | 54 | 47 | 50 |

Supplementary Table S7) *Mesoplasma* and *Spiroplasma* ASV matches to our Mollicute isolate genomes. The most abundant ASV from both the ant microbiome study [35] and the ant gut enrichment cultures (Suppl. Fig. S3) was used as a query sequences.

| | **Query** | **Reference** | **% Identity** | **Query coverage** |
|---|---|---|---|---|
| **Ant Microbiome** | *Mesoplasma* ASV | JKS002658 genome | 100 | 100 |
| | *Spiroplasma* ASV | JKS002669 genome | 100 | 100 |
| **Ant Gut Cultures** | *Mesoplasma* ASV | JKS002658 genome | 100 | 100 |
| | *Spiroplasma* ASV | JKS002669 genome | 100 | 100 |
